# Inosine Induces Stemness Features in CAR T cells and Enhances Potency

**DOI:** 10.1101/2023.04.21.537859

**Authors:** Dorota D. Klysz, Carley Fowler, Meena Malipatlolla, Lucille Stuani, Katherine A. Freitas, Stefanie Meier, Bence Daniel, Katalin Sandor, Peng Xu, Jing Huang, Louai Labanieh, Amaury Leruste, Malek Bashti, Vimal Keerthi, Janette Mata-Alcazar, Nikolaos Gkitsas, Justin A. Guerrero, Chris Fisher, Sunny Patel, Kyle Asano, Shabnum Patel, Kara L. Davis, Ansuman T. Satpathy, Steven A. Feldman, Elena Sotillo, Crystal L. Mackall

## Abstract

Adenosine (Ado) mediates immune suppression in the tumor microenvironment and exhausted CD8^+^ CAR T cells mediate Ado-induced immunosuppression through CD39/73-dependent Ado production. Knockout of CD39, CD73 or A2aR had modest effects on exhausted CAR T cells, whereas overexpression of Ado deaminase (ADA), which metabolizes Ado to inosine (INO), induced stemness features and potently enhanced functionality. Similarly, and to a greater extent, exposure of CAR T cells to INO augmented CAR T cell function and induced hallmark features of T cell stemness. INO induced a profound metabolic reprogramming, diminishing glycolysis and increasing oxidative phosphorylation, glutaminolysis and polyamine synthesis, and modulated the epigenome toward greater stemness. Clinical scale manufacturing using INO generated enhanced potency CAR T cell products meeting criteria for clinical dosing. These data identify INO as a potent modulator of T cell metabolism and epigenetic stemness programming and deliver a new enhanced potency platform for immune cell manufacturing.

**Statement of Significance:** Adenosine is well known to inhibit T cell function and substantial effort has focused on inhibiting adenosine generation and signaling. Here, we show that exhausted T cells are suppressed by adenosine, which is only modestly impacted by inhibiting adenosine generation or signaling. In contrast, metabolism of adenosine to inosine augmented T cell function and culture of T cells with inosine induced multi-level reprogramming leading to stemness and improved anti-tumor potency. We demonstrate the feasibility of introducing inosine during GMP cell manufacturing as a novel strategy to generate enhanced CAR-T cells.

## INTRODUCTION

Adenosine (Ado) is an immunosuppressive nucleoside that contributes to immune evasion in the tumor microenvironment (TME). CD39 (ecto-ATP diphosphohydrolase-1, *ENTPD1*) and CD73 (5ʹ-ectonucleotidase, *NT5E*), metabolizes ATP to Ado, which mediates immunosuppression primarily via A2aR signaling[1]. CD4^+^ Tregs co-express CD39 and CD73 and mediate immunosuppression, in part, through production of Ado[2]. CD39 is expressed by exhausted and tumor reactive CD8^+^ T cells within the TME, where it is associated with tumor progression[3–5], suggesting a role for Ado in dysfunction of exhausted T cells. Chimeric antigen receptors (CARs) link extracellular antigen binding domains to intracellular signaling domains, enabling MHC independent antigen-specific T cell reactivity[6]. CAR T cells are highly effective against refractory B cell malignancies[7, 8] but less potent against solid tumors[9–11] and exhaustion is one feature limiting the potency of CAR T cells[12, 13]. Using an *in vitro* model of human T cell exhaustion that induces hallmarks of exhaustion over 10-12 days[14, 15], we interrogated the biology of CD39 and the role of Ado in the dysfunction of exhausted human T cells.

We observed that exhausted CD8^+^CD39^+^ T cells co-express CD73, generate Ado, and mediate Ado-related suppression via A2aR. To induce Ado resistance in order to enhance potency of CAR T cells, we knocked out CD39, CD73 or A2aR, but observed only modest phenotypic and transcriptional changes. In contrast, overexpression of membrane tethered adenosine deaminase (ADA), which metabolizes Ado to inosine (INO), significantly enhanced CAR T cell fitness and altered their transcriptome. Similarly, but more potently, exposure of CAR T cells to INO induced a stemness associated gene expression program and augmented CAR T cell function.

Single cell profiling revealed that inosine reprograms the metabolome, decreasing glycolysis, but increasing glutaminolysis, polyamine synthesis and oxidative phosphorylation. These effects are associated with epigenetic reprogramming towards stemness. Clinical scale manufacturing using INO-supplemented media generated CAR T cell products meeting criteria for clinical dosing that demonstrated enhanced potency in animal models. These data identify INO as an inducer of T cell stemness and provide direct evidence that nutrient modulation can drive epigenetic changes that regulate T cell differentiation and enhance T cell fitness.

## RESULTS

### CD39 marks a subset of human exhausted T cells with diminished functionality and hallmark features of regulatory T cells

We assessed kinetics of CD39 expression on T cells expressing the GD2-targeting tonically signaling HA-CAR previously described to induce T cell exhaustion *in vitro*[14, 15]. Unlike the canonical exhaustion markers PD1, TIM3, and LAG3, which were upregulated on essentially the entire population of HA-CAR T cells by day 5 following activation and remained elevated as HA-CAR cells transition to exhaustion (**Figure 1A**), CD39 showed limited upregulation early after activation but its expression gradually increased on a subset of cells as they transitioned to exhaustion. We observed a higher frequency of CD39 expression on CAR^+^ vs. CAR^−^ cells (**Figure 1B**), and on CD8^+^ vs. CD4^+^ T cells (**Figure 1C**), and following antigen exposure, CD39^+^CD8^+^ secreted lower levels of IL-2 compared to their CD39^−^ counterparts (**Figure 1D**), but produced more TGFβ and IL-27, cytokines associated with suppressive T cells[16–18].

**Figure 1:**
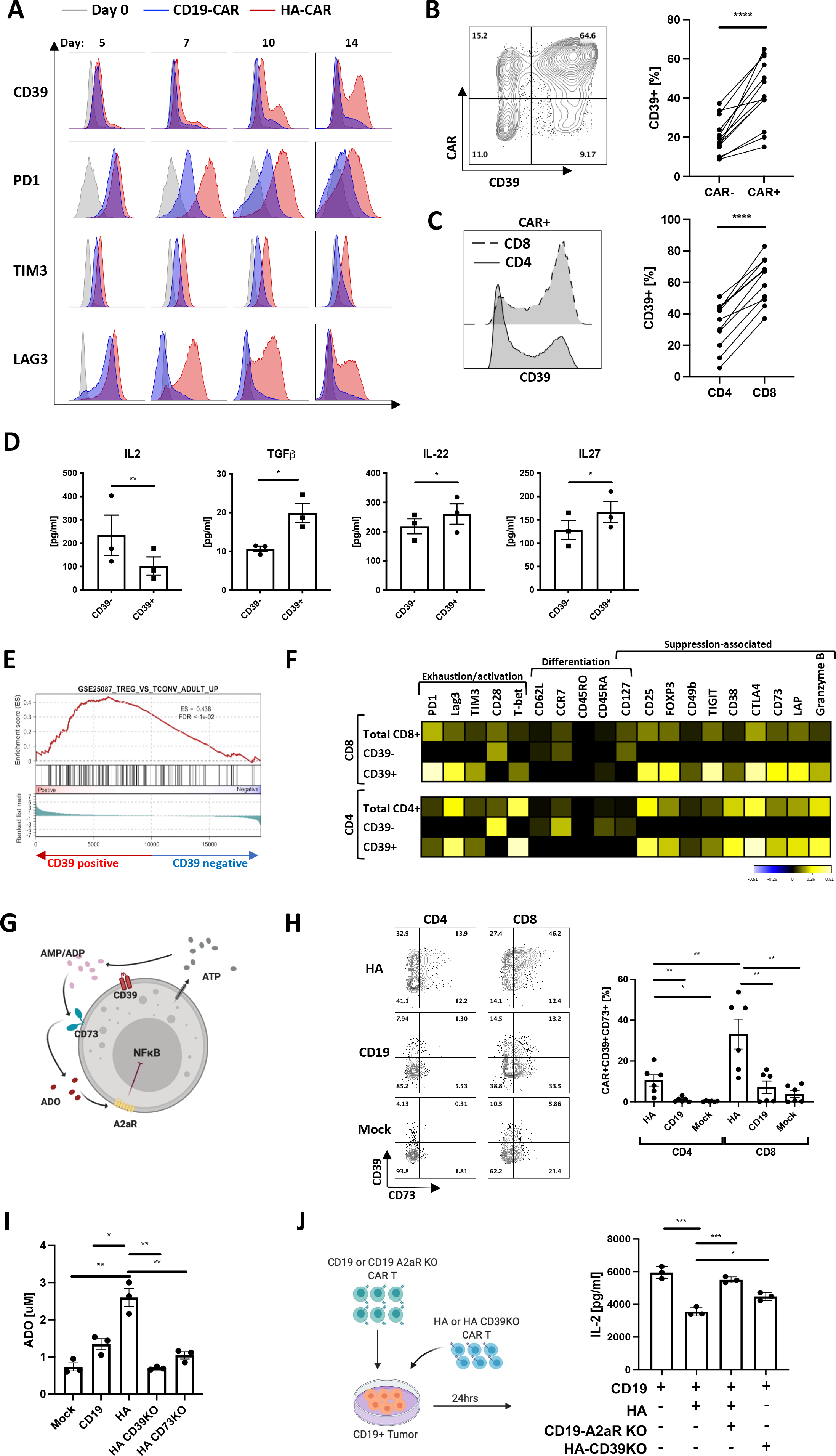
CD39 marks a subset of exhausted CAR T cells with features of regulatory T cells. A. Flow cytometric analysis of exhaustion marker expression in gated CD19.28z-CAR^+^ (CD19) (blue) and HA.28z-CAR^+^ (HA) T cells (red) at D0 (grey) and at each indicated time-point post- activation. Representative histogram shown of n = 3 donors. B. (Left) Representative flow cytometry contour plot of CD39 expression in HA-CAR^-^ vs. CAR^+^ T cells on day 10 post-activation. (Right) Percent of CD39^+^ cells in HA-CAR^_^ vs. CAR^+^ T cells from n=12 donors. P values determined by paired two-tailed t-tests. *, *P* < 0.05; **, *P* < 0.01; ***, *P* < 0.001; ****, *P* < 0.0001 C. (Left) Representative flow cytometry analysis of CD39 expression in CD8^+^ vs. CD4^+^ HA-CAR T cells 10 days post-activation. (Right) Percent of CD39^+^ cells in CD4^+^ vs. CD8^+^ HA-CAR T cells from n = 11 donors. P values determined by paired two-tailed t-tests. *, *P* < 0.05; **, *P* < 0.01; ***, *P* < 0.001; ****, *P* < 0.0001 D. Luminex analysis of cytokines secreted by CD39^+^ vs. CD39^−^CD8^+^ HA-CAR T cells sorted on day 14 and stimulated with Nalm6-GD2 for 24hrs at 1:1 E:T ratio. Data are mean ± s.e.m. of n= 3 donors. P values determined by ratio paired two-tailed t-tests. *, *P* < 0.05; **, *P* < 0.01; ***, *P* < 0.001; ****, *P* < 0.0001 E. Gene Set Enrichment Analysis (GSEA) of bulk RNA-seq collected 14 days post-activation of CD39^+^compared to CD39^−^HA-CAR T cells using the publicly available GSE25087 gene collection. F. CyTOF analysis of CD4^+^ (bottom) and CD8^+^ (top) HA-CAR T cells gated based on CD39 expression 10 days post-activation. Heat map represents expression of the indicated markers as median Arcsinh relative to the total CD4^+^ or CD8^+^ sample. Representative donor shown of n=3 donors. G. Schematic of immune suppression mediated by purinergic pathway. ATP- adenosine triphosphate; ADP- adenosine diphosphate; AMP- adenosine monophosphate; ADO- adenosine (red); A2aR- adenosine 2a receptor; CD39- ecto-ATP diphosphohydrolase-1; CD73- 5ʹ-ectonucleotidase; NF- κB- Nuclear Factor kappa B. H. (Left) Representative flow cytometry contour plots showing expression of CD39 and CD73 in gated CD4^+^ or CD8^+^ HA-, CD19-CAR or mock T cells 14 days post-activation. (Right) Percent of CD39^+^CD73^+^ T cells in CD4^+^ vs. CD8^+^ by day 14 HA-, CD19-CAR or mock T cells in 6 donors. P values determined by paired two-tailed t-tests. *, *P* < 0.05; **, *P* < 0.01; ***, *P* < 0.001; ****, *P* < 0.0001 I. Concentration of adenosine (ADO) produced by mock or CAR-expressing T cells with or without CD39 or CD73 spiked with 20uM of ATP at day 17 post-activation. Representative data from n= 3-5 donors. Average purity of the knock-out cells was > 90%. P values determined by unpaired two-tailed t-tests. *, *P* < 0.05; **, *P* < 0.01; ***, *P* < 0.001; ****, *P* < 0.0001 J. (Left) schematic of experimental design of immunosuppression co-culture assays. (Right) Control or A2aR KO CD19.bbz-CAR T cells were activated with Nalm6 at 1:1 ratio in the presence of HA- or CD39KO HA-CAR T cells. IL-2 was measured 24hrs post-stimulation. Data are mean ± s.e.m. from triplicate wells. P values determined by unpaired two-tailed t-tests. Representative result of two independent experiments. *, *P* < 0.05; **, *P* < 0.01; ***, *P* < 0.001; ****, *P* < 0.0001

CD39^+^ CAR T cells also demonstrated distinct transcriptomic profiles compared to CD39^−^ subsets, with higher levels of exhaustion/effector-related genes (*NR4A1, NR4A2, NR4A3* and *KLRD1*)[19], increased Treg-associated genes (*FOXP3, TIGIT, IL2RA, RXRA)*[20], and lower stemness-associated genes such as *TCF7, TCF4, SELL, LEF1* and *IL-7R* (**Figure S1A**). These findings were confirmed by Gene Set Enrichment Analysis (GSEA), which demonstrated that CD39^+^CD8^+^ and CD4^+^ CAR T cells resemble exhausted T cells [14, 21] and CD4^+^ regulatory T cells (**Figure 1E, S1B**), whereas CD39^−^ CAR T cells resembled non-exhausted CD19-CAR T cells. Mass cytometry confirmed higher levels of exhaustion-associated (TIM3, PD1, LAG3) and suppression-associated (Foxp3, TIGIT, CD49b, LAP, CD73, CTLA4) proteins in CD39^+^ vs. CD39^−^ HA-CAR T cells (**Figure 1F**).

The CD39^+^ subset in this model could represent expansion of a preexisting CD39^+^ population or CD39 induction in response to tonic signaling. To distinguish between these possibilities, we sorted CD39^−^ HA-CAR T cells and cultured them ± dasatinib, a tyrosine kinase inhibitor that blocks CAR signaling, then assessed CD39 expression[15, 22] (**Figure S1C**). CD39^−^ HA-CAR cells rapidly re-expressed CD39, and this was prevented by dasatinib, confirming a requirement for CAR signaling to CD39 upregulate in this model. TGFβ contributes to CD39 upregulation in regulatory T cells[23], but neutralizing anti-TGFβ antibody did not affect the frequency of CD39^+^ cells, nor expression levels of CD39 on HA-CAR T cells in this system. Together, these data demonstrate that CD39 expression is induced by chronic T cell activation and marks a highly dysfunctional exhausted T cell subset with hallmark features of regulatory T cells.

### A high fraction of exhausted CD39^+^ T cells co-express CD73, generate Ado and mediate immunosuppression through A2aR

CD39 metabolizes extracellular ATP (eATP) to ADP/AMP and CD73 metabolizes ADP/AMP to Ado, which signals via A2aR to induce 3’,5’-cyclic Ado monophosphate (cAMP) and can inhibit NF-κB[24] (**Figure 1G**). We observed high frequencies of CD39^+^CD73^+^ co-expression in HA-CAR T cells, whereas dual CD39^+^CD73^+^ expression was rare in CD19 or mock non-transduced cells (**Figure 1H**). The frequency of CD39^+^CD73^+^ cells was significantly higher in CD8^+^ vs.CD4^+^ HA-CAR T cells. CD39 and CD73 are enzymatically active on HA-CAR T cells since they induced greater extracellular ATP (eATP) hydrolysis (**Figure S1D**) and generated more Ado than mock or control CD19-CAR T cells (**Figure 1I**). Knockout of CD39 or CD73 from HA-CAR T cells (HA- CD39KO and HA-CD73KO) using CRISPR/Cas9, confirmed the requirement for CD39 to induce eATP hydrolysis to ADP/AMP, and for CD73 to generate Ado in this system.

To test whether CAR T cells are susceptible to Ado-mediated suppression, we activated HA- or CD19-CAR T cells with antigen, in the presence or absence of the Ado receptor agonist 5′-(N-ethylcarboxamido) adenosine (NECA). NECA reduced antigen induced IL-2 and IFNγ production (**Figure S1E**), but this inhibition was prevented by CPI-444, a selective A2aR competitive antagonist (iA2aR)[25]. To assess whether CD8^+^ HA-CAR T cells mediate immune suppression and whether this is mediated by Ado, we stimulated CD19-CAR T cells in the presence of bulk HA-CAR T cells or sorted CD8^+^ HA-CAR T cells and observed decreased antigen-induced IL-2 production by CD19-CAR T cells (**Figure S1F)**. CD39 expression was required by the suppressive population and A2aR expression in the responder since CD39KO HA-CAR T cells mediated less suppression while CD19-CAR T cells with A2aR KO restored IL-2 secretion to the level of control CD19-CAR T cells (**Figure 1J**). Together, data demonstrate that CD39 marks a highly dysfunctional exhausted T cell subset that bear hallmark features of regulatory T cells and mediate immunosuppression via Ado.

### Overexpression of Ado deaminase (ADA-OE) induces transcriptomic and phenotypic features of T cell stemness

With the goal of developing a cell intrinsic approach to prevent Ado mediated immunosuppression, we knocked out CD39, CD73 or A2aR on HA-CAR T cells and we also incorporated an experimental group with membrane tethered overexpression of adenosine deaminase (ADA-OE) (**Figure 2A, S2A**), reasoning that ADA will catabolize Ado to INO, and reduce Ado mediated immunosuppression. Neither ADA-OE nor KO of CD39, CD73 or A2aR significantly impacted CAR expression (**Figure S2B**). ADA-OE induced the most significant changes in gene expression compared to control (**Figure 2B**), upregulating genes associated with stemness (*TCF7, IL-7R, SELL*) and glutamine transport (*SLC1A5*) and downregulating genes associated with effector function (*GZMB, IFNγ*, *IL-3, IL-5, TNFSF4* (OX40) and *TNFSF11* (RANKL) (**Figure 2C**).

**Figure 2:**
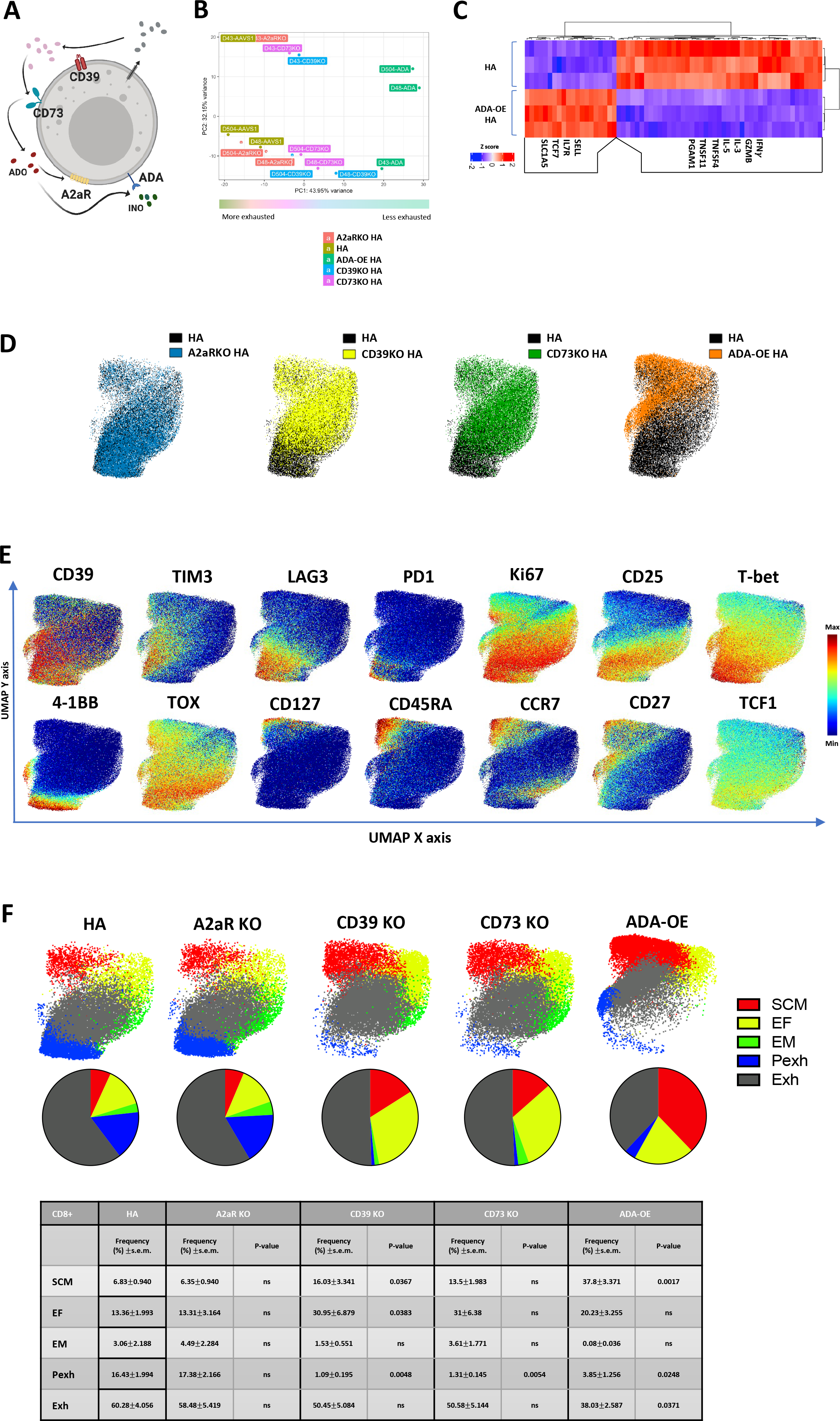
Overexpression of Adenosine deaminase (ADA-OE) increases features of stemness and decreases terminal differentiation. A. Schematic of immune suppression mediated by purinergic pathway. ADO- adenosine (red); A2aR- adenosine 2a receptor; CD39- ecto-ATP diphosphohydrolase-1; CD73- 5ʹ-ectonucleotidase; INO- inosine (green); ADA- adenosine deaminase. B. Principal component analysis (PCA) of bulk RNA-seq from HA-CAR T cells overexpressing Ado deaminase (ADA-OE) or deficient for CD39 or CD73 15 days post-activation. Data from n=3 donors. C. Gene expression of the top differentially expressed genes between HA-ADA-OE vs. HA-AAVS1- KO (HA) CAR T cells identified by DESeq2. C-E. UMAP analysis of day 15 CD8^+^ HA-CAR cells engineered as in (A). Expression of 26 markers was analyzed by CyTOF. 5,000 or maximum of CD8^+^ CAR T cells from each donor (n=4) were combined and colored by (C) genotype, (D) marker intensity or (E) subpopulation defined by FlowSOM algorithm. Pie charts show population frequencies defined using FlowSOM for each condition. SCM; stem cell memory, EF; effector, EM; effector memory, Pexh; progenitor exhausted, Exh, exhausted. Table shows the average frequencies from n=4 donors and corresponding p values determined by paired two-tailed t-tests.

Consistent with this, GSEA analysis of ADA-OE HA-CAR T cells showed a negative enrichment score with gene sets associated with T cell exhaustion[14, 21] and a positive enrichment score with gene sets of non-exhausted CD19-CAR T cells and CD19-CAR T cells associated with long-term complete response[14, 26] (**Figure S2C**).

To assess whether the transcriptomic changes observed translated to protein expression, we used mass cytometry to evaluate a panel of 26 surface markers and transcription factors (TFs) in control vs. CD39KO, CD73KO, A2aR KO and ADA-OE HA-CAR T cells (**Figure 2D**). Consistent with transcriptomic analysis, ADA-OE HA-CAR T cells were most significantly different from controls based upon higher expression of stemness markers (CD27, CD45RA and CD127), and lower expression of exhaustion (LAG3, PD1, T-bet, TOX) and activation (CD25 and 4-1BB) markers (**Figure 2E, S2D**). FlowSOM analysis[27] of CD8^+^ ADA-OE vs. control HA-CAR T cells revealed significantly increased frequencies of stem-like memory T cells (SCM) (CCR7^hi^CD127^hi^CD95^hi^CD62L^hi^), and decreased progenitor exhaustion (Pexh) (TCF1^+^TOX^+^T-bet^+^CD45RO^+^Ki67^+^) and terminally differentiated exhausted-like (Exh) populations[28, 29] (**Figure 2F**). Similar findings were seen in CD4^+^ ADA-OE cells compared to control HA-CAR T cells (**Figure S2E-F)**. CD39KO and CD73KO cells manifested an intermediate phenotype with reduced fractions of Pexh cells, no change in frequency of Exh cells compared to control, but increased fractions of stem-like memory T cells and effector subsets (EF) (**Figure 2F**). Together, the data demonstrate that ADA-OE, and to a lesser extent CD39KO or CD73KO but not A2aR KO, induces transcriptomic and phenotypic features of increased stemness, raising the prospect that ADA-OE could be inducing effects beyond inhibition of Ado signaling. Since ADA-OE is predicted to generate INO, and T cells can use inosine as an alternative energy source to glucose for metabolic processes[30], we hypothesized that INO could be responsible for the findings.

### INO increases stemness features, enhances T cell functionality, and abrogates immune suppression in chronically activated CAR T cells independently of the presence of glucose

To test the hypothesis that INO could directly modulate stemness in chronically activated CAR T cells, we activated, transduced and cultured cells in standard RPMI containing 11mM of glucose (unchanged from previous experiments) and, on day 4, split cultures into standard RPMI or RPMI containing 11 mM of INO, instead of glucose (**Figure 3A**). Transcriptomic analysis on day 14 demonstrated that HA-INO-CAR T cells manifested differential expression of >3,000 gene transcripts compared to cells expanded in control media, with INO exposure responsible for 41.3% of variance observed in principal component analysis (PCA) (**Figure 3B**). GSEA analysis demonstrated that HA-INO-CAR T cells upregulated transcriptional programs associated with stem cell-like memory population (**Figure 3C**) and downregulated genes involved in the purinergic pathway CD73 (*NT5E*), A2aR (*ADORA2A*) and FOXP3 (**Figure 3D**). Flow cytometry following INO exposure demonstrated higher levels of CD62L (**Figure 3E**) and decreased frequencies of CD39^+^CD73^+^ cells, largely due to reduced CD73 expression (**Figure S3A**). Increased protein expression of stemness markers (CD45RO, CCR7, and CD127) in CD8^+^ (**Figure S3C**) and CD4^+^ (**Figure S3B-C**) HA-INO-CAR T cells was confirmed by CyTOF analysis.

**Figure 3:**
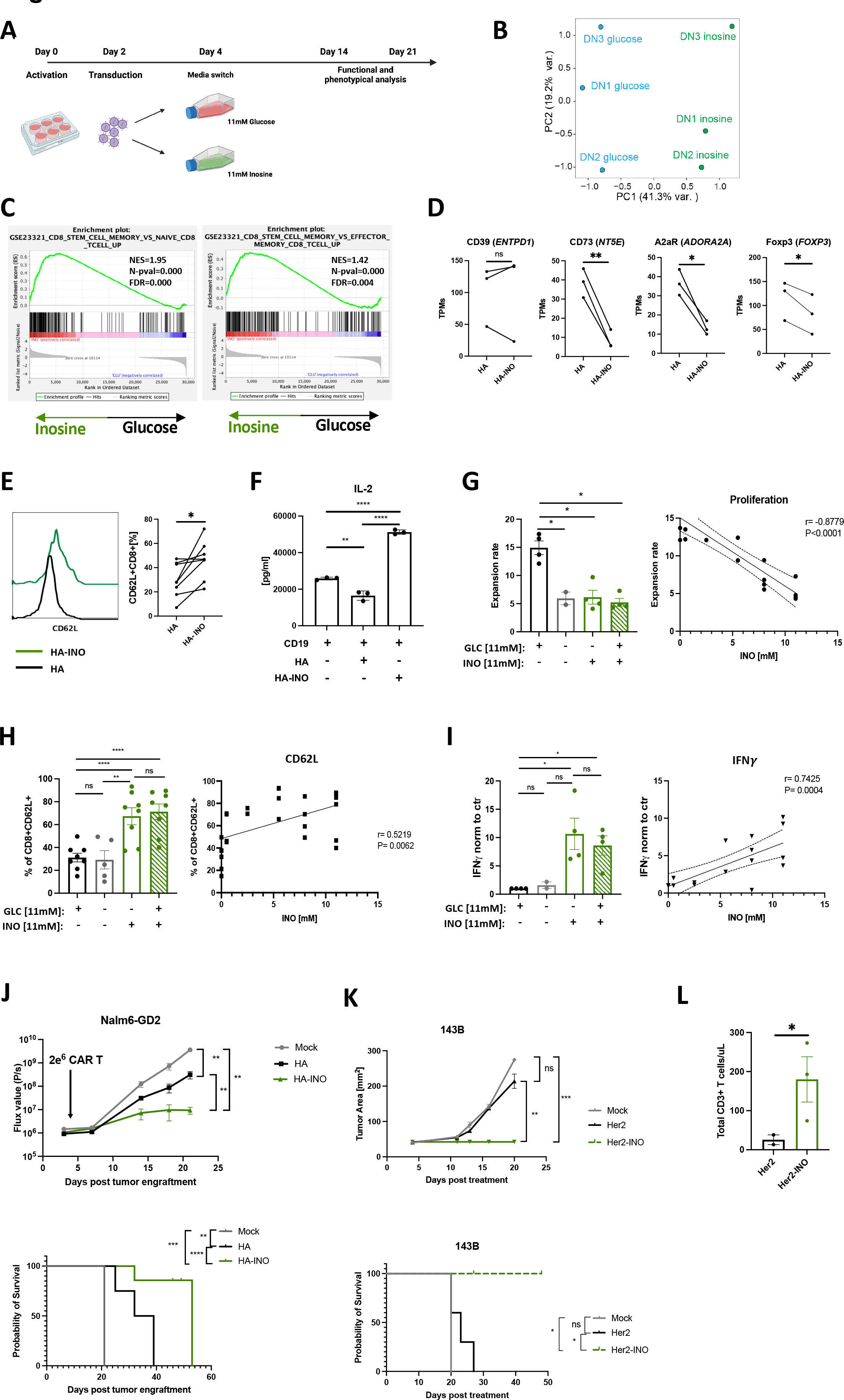
Inosine (INO) induces stemness and augments CAR T cell function in vivo. A. Schematic of the experimental design. B. PCA of bulk RNA-seq from HA-CAR T cells expanded in glucose vs. inosine containing media for 10 days. C. GSEA of HA-CAR T cells expanded as in (B) using the GSE23321 gene collection. D. Levels of expression of indicated purinergic pathway gene transcripts from bulk RNA samples shown in (B) Data are mean+/- s.e.m. from n=3 donors. P values determined by paired two-tailed t-tests. *, *P* < 0.05; **, *P* < 0.01; ***, *P* < 0.001; ****, *P* < 0.0001 E. Flow cytometry analysis of CD62L expression by CD8^+^ HA-CAR T cells at day 21 post-activation. Histograms of representative donor (left) and pooled data from n=8 donors shown (right). P values determined by paired two-tailed t-tests. *, *P* < 0.05; **, *P* < 0.01; ***, *P* < 0.001; ****, *P* < 0.0001 F. IL-2 release after 24h of CD19.4-1BBz-CAR T cells stimulated with Nalm6 tumor line alone or in co-culture with HA- or HA-INO-CAR T cells at 1:1:1 ratio. Data are mean ± SD from triplicate wells. N=1 donor. *, *P* < 0.05; **, *P* < 0.01; ***, *P* < 0.001; ****, *P* < 0.0001 G. (Left) Cell expansion of HA-CAR T cells grown in culture media containing indicated concentration of glucose and/or inosine from day 4 to day 10 post-activation. Data are mean ± s.e.m. of n = 2-4 donors. P values determined by paired two-tailed t-tests. *, *P* < 0.05; **, *P* < 0.01; ***, *P* < 0.001; ****, *P* < 0.0001 (Right) Expansion of HA-CAR T cells grown in RPMI containing 11mM of glucose and increasing concentrations of inosine for 6 days. Scatter plot showing correlation between ratio of expansion (y-axis) and inosine concentration (x-axis). Pearson correlation, r, with P values determined by paired two-tailed t-tests. N= 2-4 donors from independent experiments. H. (Left) Percent of CD62L^+^CD8^+^ HA-CAR T cells grown in culture media containing indicated concentration of glucose and/or inosine from day 4 to day 21 and measured by flow cytometry. Data are mean ± s.e.m. of n = 5-8 donors. P values determined by paired two-tailed t-tests. *, *P* < 0.05; **, *P* < 0.01; ***, *P* < 0.001; ****, *P* < 0.0001 (Right) HA-CAR T cells were grown in RPMI containing 11mM of glucose and increasing concentrations of inosine for 17 days. Scatter plot showing correlation between CD62L frequency (y-axis) and inosine concentration (x-axis). Pearson correlation, r, with P values determined by paired two-tailed t-tests. N= 2-6 donors from independent experiments. I. (Left) Fold increase of IFNγ release after 24hrs of co-culture with Nalm6-GD2 by D14 HA-CAR T cells grown in culture media containing various concentrations of glucose and/or inosine and normalized to CAR T cells grown in control media. Data are mean ± s.e.m. of n = 2-4 donors. P values determined by paired two-tailed t-tests. *, *P* < 0.05; **, *P* < 0.01; ***, *P* < 0.001; ****, *P* < 0.0001 (Right) HA-CAR T cells were grown in RPMI containing 11mM of glucose and increasing concentrations of inosine for 10 days, and stimulated with Nalm6-GD2 for 24hrs. Scatter plot showing correlation between IFNγ secretion (y-axis) and inosine concentration (x-axis). Pearson correlation, r, with P values determined by paired two-tailed t-tests. N= 2-4 donors from independent experiments. J. NSG mice were injected intravenously with 1 × 10^6^ Nalm6 leukemia cells. Three days later 2 × 10^6^ of mock or HA-CAR T cells manufactured in the presence of inosine- or glucose-containing media from day 6 to 10 post-activation, were given intravenously. (Top) Tumor progression was monitored using bioluminescent imaging. Data are mean ± s.e.m. of n = 5 mice per group. P values determined at day 21 by using Mann-Whitney test. (Bottom) Survival curves were compared using the log-rank Mantel–Cox test. *, *P* < 0.05; **, *P* < 0.01; ***, *P* < 0.001; ****, *P* < 0.0001 K. NSG mice were inoculated with 1 × 10^6^ 143B osteosarcoma cells via intramuscular injection. At D4 post-engraftment 1 × 10^7^ mock or Her2.bbz-CAR T cells cultured in glucose- or inosine- containing media were given intravenously. (Top) 143B tumor growth monitored by caliper measurements. P values determined by unpaired two-tailed t-test with Welch’s correction at day 20. (Bottom) Long-term survival of CAR-treated mice. Data are mean ± s.e.m. of n = 3-5 mice per group. Survival curves were compared using the log-rank Mantel–Cox test. *, *P* < 0.05; **, *P* < 0.01; ***, *P* < 0.001; ****, *P* < 0.0001 L. Concentration of total CD3^+^ T cells detected in blood of mice at day 17 post-CAR T cell injection. P values determined by unpaired two-tailed t-test with Welch’s correction. *, *P* < 0.05; **, *P* < 0.01; ***, *P* < 0.001; ****, *P* < 0.0001

Functional studies following challenge with tumor cell lines expressing high (Nalm6-GD2) or low (143b) levels of GD2 (**Figure S3D**) revealed increased cytokine production and cytotoxicity by both ADA-OE and HA-INO-CAR vs. control HA-CAR T cells (**Figure S3E-F**), with HA-INO- CAR T cells showing highest potency. We also assessed suppressive effects of HA-INO-CAR T cells and observed that CD19-CAR T cells activated in the presence of control HA-CAR T cells exhibited significantly decreased secretion of IL-2, whereas those activated in the presence of HA- INO-CAR T cells demonstrated increased IL-2 secretion compared to control (**Figure 3F**).

Next, we tested if the changes observed could be attributed to INO addition or glucose deprivation, since transient glucose restriction has been demonstrated to augment T cell function [31]. HA-CAR T cells were cultured from day 4 post-activation in control media containing glucose, media without glucose or inosine, inosine only as described above (INO) or media containing both inosine and glucose (**Figure S3G**). On day 10 we observed that INO significantly reduced expansion in a dose-dependent manner, whether or not glucose was present (**Figure 3G**). INO also significantly enhanced CD62L expression, and the frequency of central memory CD8^+^ HA-CAR T cells, independently of glucose (**Figure 3H, S3H**), and INO significantly enhanced antigen induced IFNγ secretion in the presence or absence of glucose (**Figure 3I**). Together these data demonstrate INO restrains proliferation of chronically activated T cells, induces a more stem-like phenotype, endows enhanced functionality and eliminates of suppressive activity and that these effects are not replicated by glucose deprivation.

### CAR T cells cultured in inosine mediate superior tumor control *in vivo*

To test if INO induced phenotypic and functional changes in CAR T cells translate into improved antitumor activity, we conducted an *in vivo* “CAR stress test” using low numbers of CAR T cells in the Nalm6-GD2 leukemia model administered to NSG mice. HA-INO-CAR T cells significantly delayed tumor growth and increased overall survival compared to HA-CAR T cells (**Figure 3J, S4A-B**). Tumor cells expressing target antigen below the activation threshold are a major cause of resistance to CAR T cells[32], thus we assessed whether CD19-INO-CAR T cells manifested enhanced capacity to control antigen low tumor cells. CD19-INO-CAR T cells exhibited increased cytokine production *in vitro* compared to control and ADA-OE CD19-CARs (**Figure S4C**) and mediated markedly increased anti-tumor activity *in vivo* in a CAR stress test against CD19 leukemia (**Figure S4D**). Next, we tested if INO-CAR T cells manifested enhanced functionality against solid tumors where exhaustion and Ado mediated immunosuppression are known to be a major cause of therapeutic failure. We assessed the activity of Her2-CAR T cells, a CAR which has been used by others to study Ado-related immunosuppression[33], in the 143B osteosarcoma mouse model (**Figure 3K, S4E**). Tumors treated with mock or Her2-CAR T cells grew at the same rate, whereas Her2-INO-CAR T cells prevented tumor growth and prolonged survival. We further observed 10 times more circulating T cells in the blood of mice 17 days after receiving the Her2- INO-CAR vs. control Her2-CAR T cells (**Figure 3L**). These data demonstrate that INO improves antitumor potency of CAR T cells prone to early exhaustion as well as those, which are not prone to exhaustion, such as CD19-CAR T cells and that INO enhanced CAR T cell persistence.

### INO modifies metabolic programming of chronically activated CAR T cells

To better understand the mechanisms responsible for these results, we conducted DAVID gene annotation enrichment analysis using KEGG pathways and GO terms (**Figure 4A**). We observed substantial evidence for metabolic reprogramming, with downregulation of genes associated with glycolysis and glucose driven metabolism (*PFKP*, *ALDOA*, *GAPDH*, *PGK*, *ENO1*), and increased expression of genes associated with glutaminolysis and polyamine synthesis (**Figure 4B-C**). INO increased expression of ornithine decarboxylase (ODC), the proximal, rate limiting enzyme for putrescine synthesis and spermidine synthase, which generates spermidine and eIF5A, important mitochondria activity regulators. Previous work has implicated polyamines in epigenetic regulation of Th1 differentiation, raising the prospect that increased polyamines may drive the epigenetic stemness programming induced by INO exposure[34]. These metabolic effects were also associated with diminished expression of genes associated with TCR signaling consistent with decreased effector differentiation.

**Figure 4:**
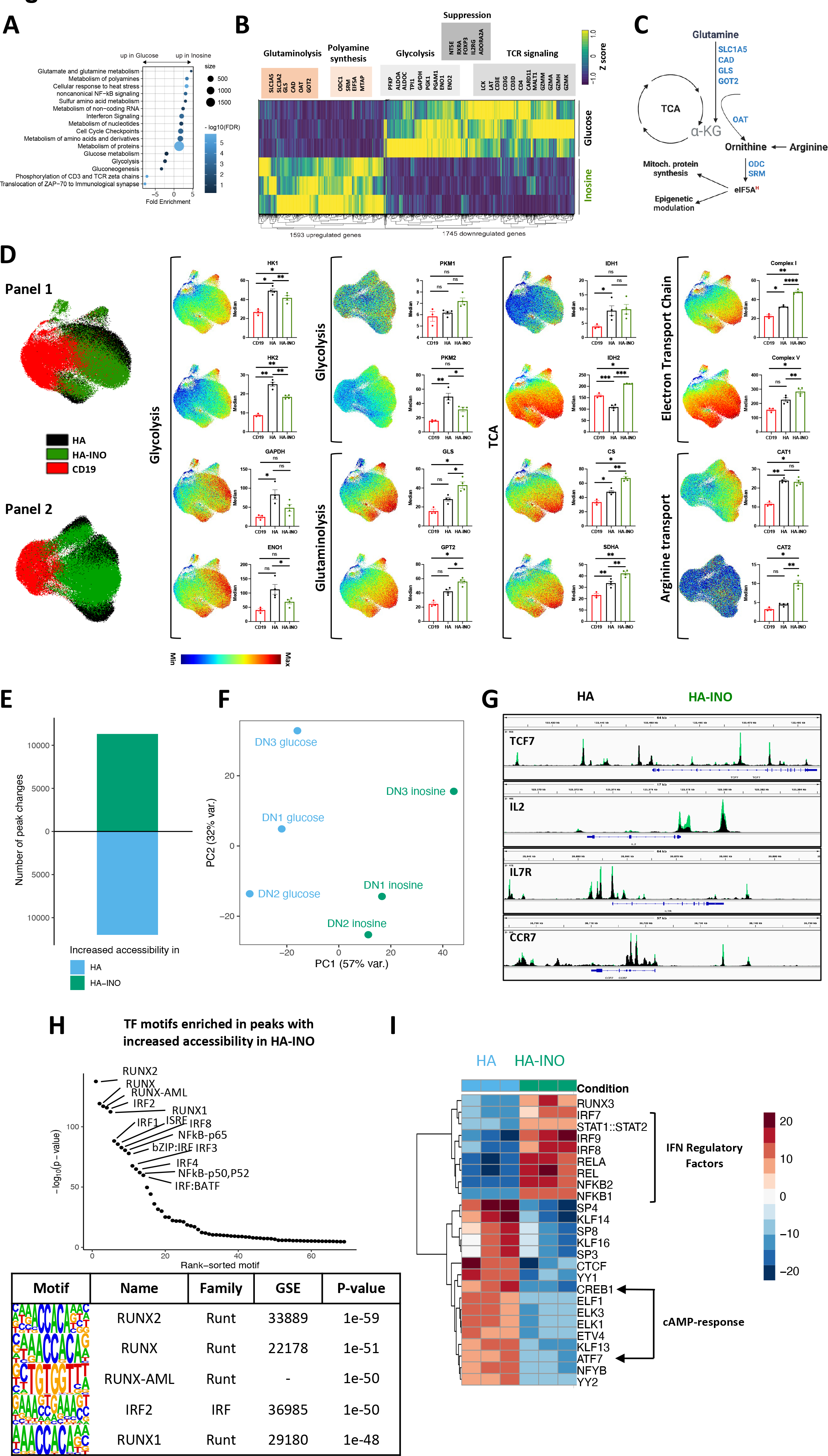
Exhausted CAR T cells expanded in inosine undergo metabolic, epigenetic and phenotypic changes. A. Pathway Enrichment Analysis of day 14 CAR-T cells shown in (3B) using the Reactome pathway collection and DAVID algorithm. Fold enrichment, number of genes represented, and FDR are shown. B. Heat map of differentially expressed genes in HA vs. HA-INO CAR-T cells shown in (3B) (padj<0.01). C. The glutaminolytic pathway and polyamines synthesis pathway, with the metabolic genes upregulated in (B) highlighted in blue. D. UMAP analysis of CD8^+^ CD19.28z-, HA- or HA-INO-CAR T cells 14 days post-activation. Expression of 28 markers (Panel 1) or 24 markers (Panel 2) was analyzed by CyTOF. 9,000 of CD8^+^ CAR T cells from each condition and donor (n=3-4) were combined and colored by marker intensity. Graphs represent median expression of indicated markers expressed by CAR^+^CD8^+^ T cells. P values determined by paired two-tailed t-tests. *, *P* < 0.05; **, *P* < 0.01; ***, *P* < 0.001; ****, *P* < 0.0001 E. Global chromatin accessibility profile of D14 CD8^+^ HA- and HA-INO CAR T cells from three healthy individuals determined by ATACseq (p-value < 0.001 and log2 FC > 1). F. PCA of ATACseq chromatin accessibility from HA-CAR T cells cultivated in glucose or inosine. G. Overlayed accessibility profiles in HA (black) and HA-INO-CAR T cells (green) in the *TCF7, IL2, IL7R* and *CCR7* loci at day 14 post-activation. Concatenated samples from n=3 donors. H. Top transcription factor motifs enrichment in HA-INO vs. HA- T cells ranked by HOMER analysis. I. Top 25 transcription factor motif deviation scores in HA-INO versus HA- by chromVAR analysis.

To assess whether protein expression also demonstrated evidence for widespread metabolic reprogramming, we used mass cytometry to perform single cell metabolic regulome profiling[35] using two panels of antibodies targeting metabolic enzymes and nutrient transporters (**Figure 4D**). Compared to HA-CAR T cells, the proteome of HA-INO-CAR T cells demonstrated reduced levels of enzymes associated with glycolysis, including reduced hexokinases (HK1/2), glyceraldehyde 3-phosphate dehydrogenase (GAPDH), enolase 1 (ENO1) and PKM2. Importantly however, glycolysis was not completely inhibited, since glycolytic enzymes were present at similar levels in HA-INO-CAR T cells and control CD19-CAR T cells, consistent with previous evidence that INO is capable of providing a carbon source for glycolysis[30]. INO cultured cells also expressed higher levels of enzymes associated with glutaminolysis (GLS, GPT2), and showed evidence for increased TCA activity (IDH1, IDH2, CS, SDHA), consistent with INO induced glutamine catabolism driving TCA through anapleurosis (**Figure 4D**). CAT2, a high efficiency transporter of the polyamine precursor-arginine[36] also increased following INO culture, providing further evidence in support of increased polyamine synthesis (**Figure 4D**). Proteins of complex I and complex V of the electron transport chain were also elevated, consistent with higher oxidative phosphorylation. Together, phenotypic, transcriptomic and proteomic profiling demonstrate that INO induced stemness programming augmented functionality is associated with profound metabolic reprogramming, characterized by diminished glycolysis, increased glutaminolysis and polyamine synthesis, and increased TCA and mitochondria activity.

### Inosine modulates the epigenetic landscape toward stemness

T cell exhaustion is associated with widespread epigenetic changes[14, 15, 37] and metabolic pathways are increasingly implicated in modulating the epigenome of T cells[34, 38]. To assess whether INO induced functional and metabolic reprogramming was associated with epigenetic reprogramming of exhausted T cells, we used ATACseq to compare the chromatin landscape of CD8^+^ HA- and HA-INO-CAR T cells. INO induced widespread changes in chromatin accessibility spanning more than 10,000 genomic regions (PC1 variance 57%) (**Figure 4E-F**). Enhanced chromatin accessibility in HA-INO-CAR T cells was evident in regulatory regions associated with IL-2 and several stemness genes (*TCF7, IL-7R* and *CCR7*) (**Figure 4G**). Using ChromVAR and HOMER transcription factor binding motif enrichment analysis we discovered that INO increased accessibility of RUNT (RUNX1/2), IRF (IRF1/2/3/4/8, ISRE) and RHD (NFKB1/2, REL(A)) family TF binding motifs, all which are associated with memory differentiation[39] (**Figure 4H**). In contrast, accessibility of cAMP-response element binding proteins CREB1 and ATF7 were decreased in INO-HA compared to HA-CAR T cells, raising the prospect that INO exposed cells are less susceptible to Ado-mediated suppression (**Figure 4I**). Together, the data demonstrate that INO induces metabolic and epigenetic programming in T cells that drive enhanced functionality, induces features of stemness and diminishes susceptibility to Ado mediated immune suppression.

### Large scale manufacturing of clinical grade GD2-CAR T cells using inosine-containing media improves CAR product quality

We next tested whether it is feasible to manufacture GD2-CAR T cells, which have demonstrated anti-tumor activity in diffuse midline gliomas (DMGs)[40] using INO in a GMP compliant process at clinical scale. A total of 2.5 x10^8^ enriched CD4^+^ and CD8^+^ T cells from a healthy donor were activated with anti-CD3/CD28-coated beads and transduced using clinical grade GD2-CAR vector at an MOI of 10 (**Figure 5A**). On day 3, T cells were maintained in standard RPMI media containing glucose or switched to inosine-containing RPMI without glucose in a semi-closed G- Rex culture platform. On day 7, there was no difference in transduction efficiency, CD4/CD8 ratio or viability between control and INO CAR^+^ T cells (**Figure 5A**). The total cell yield in INO cultures was reduced compared to control cultures as predicted (238.5x10^6^±145.8 vs. 387.7x10^6^±165 s.e.m., respectively), however, these yields were sufficient to formulate clinical doses of 1 x10^6^ CAR+ T cells/kg body weight (+/- 20%) (**Figure 5B**). Consistent with the results from the small-scale experiments, we observed a higher frequency of naive and central memory GD2-CAR T cells in INO cultures (**Figure 5C**). When stimulated with 143b (low GD2 expression), mg63.3 or Nalm6-GD2 (high GD2 expression) tumor cells, GD2-INO-CAR T cells secreted significantly more IL-2 and IFNγ (**Figure 5D**), mediated significantly greater antitumor effects *in vivo* (**Figure 5E**), and exhibited greater persistence when compared to control GD2- CAR T cells (**Figure 5F**).

**Figure 5.**
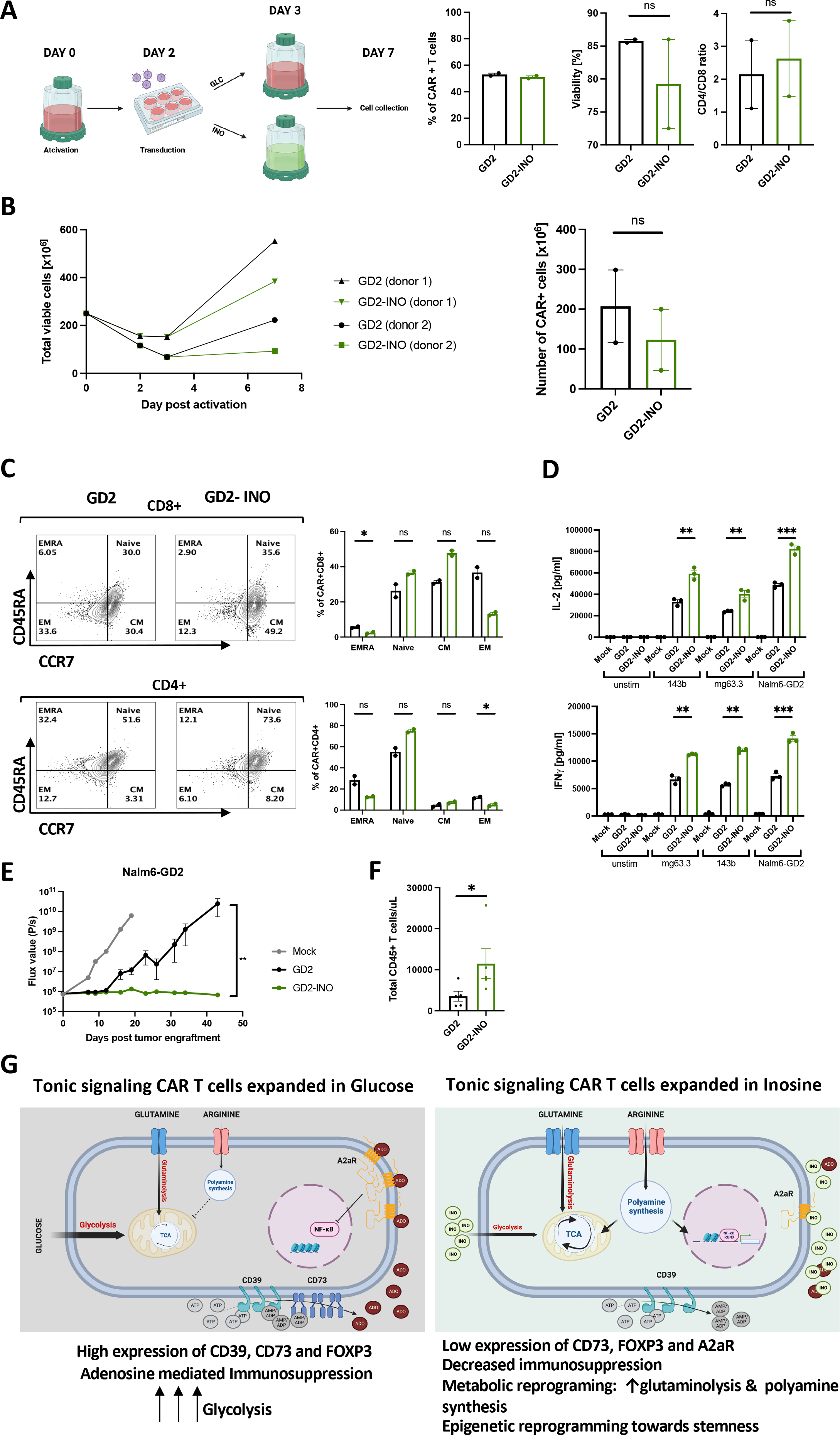
Clinical scale manufacturing of clinical grade GD2-CAR T cells using media containing inosine, improves CAR product quality. A. Schematic of large-scale manufacturing of GD2.bbz-CAR^+^ T cells in a semi-closed G-Rex system in glucose- or inosine-containing media. 7 days post-activation, transduction efficiency, percentage of viable cells, and CD4/CD8 ratio of GD2^+^ cells were assessed by flow cytometry. Data are mean+/- s.e.m. from n=2 donors. P values determined by paired two-tailed t-tests. *, *P* < 0.05; **, *P* < 0.01; ***, *P* < 0.001; ****, *P* < 0.0001 B. (Left) Expansion of total viable T cells manufactured in control (black) vs. inosine (green)- containing media and (Right) corresponding number of CAR+ cells at day 7. Data are mean+/- s.e.m. from n=2 donors. P values determined by paired two-tailed t-tests. *, *P* < 0.05; **, *P* < 0.01; ***, *P* < 0.001; ****, *P* < 0.0001 C. Flow cytometry analysis of the relative frequency of terminally differentiated effector (EMRA; CD45RA^+^CCR7^−^), naïve (CD45RA^+^CCR7^+^), central memory (CM; CD45RA^−^CCR7^+^), and effector memory (EM; CD45RA^−^CCR7^−^) in CD8^+^ (top) and CD4^+^ (bottom) control or inosine- grown GD2-CAR T cells. Representative donor shown. N=2 donors. *, *P* < 0.05; **, *P* < 0.01; ***, *P* < 0.001; ****, *P* < 0.0001 D. IL-2 and IFNγ secretion by CAR T cells stimulated for 24hrs with 143b, mg63.3 or Nalm6-GD2 tumor lines. Error bars represent mean ± SD of triplicate wells from one representative donor (n=2 donors). P values determined by unpaired two-tailed t-tests. *, *P* < 0.05; **, *P* < 0.01; ***, *P* < 0.001; ****, *P* < 0.0001 E. NSG mice were injected with 1 × 10^6^ Nalm6 leukemia cells. On D3 post-tumor injection, 2 × 10^6^ of mock or GD2.bbz-CAR T cells manufactured in the presence of inosine or in regular media were transferred intravenously. Tumor growth was monitored by bioluminescent imaging. Data are mean ± s.e.m. of n=5 mice per group. P values determined at day 43 by using Mann-Whitney test. Representative results of two independent experiments shown. *, *P* < 0.05; **, *P* < 0.01; ***, *P* < 0.001; ****, *P* < 0.0001 F. Concentration of CD45^+^ T cells detected in blood of mice at D33 post-CAR T cell injection. P values determined by unpaired two-tailed t-test with Welch’s correction. Results from one experiment shown. *, *P* < 0.05; **, *P* < 0.01; ***, *P* < 0.001; ****, *P* < 0.0001 G. Proposed model showing metabolic and epigenetic differences between tonic signaling CAR-T cells expanded in glucose- vs. inosine-containing culture media. Reprograming is due to an increase in glutaminolysis and polyamine synthesis and downregulation of CD73, FOXP3, A2aR and c-AMP response transcription factors. This results in an increase of stemness and a reduction of the immunosuppressive phenotype.

Finally, to test compatibility of adding INO to GMP-grade culture media that contains glucose, we manufactured GD2-CAR T cells in TexMACS^TM^ with increasing concentrations of INO (**Figure S5A**). We observed significantly decreased cell proliferation (**Figure S5B**) but no changes in CAR T cell viability. INO increased CD62L expression compared to control cells (**Figure S5C**) and we observed a dose-dependent increase in secretion of IL-2 and IFNγ when cells were challenged with 143b cells (**Figure S5D**). While INO concentrations between 5-11mM mediated similar anti-proliferative effects and CD62L induction, we observed a trend toward greater antigen induced cytokine production with INO concentrations of 11mM (**Figure S5D**). Together, these data demonstrate that CAR T cells can be manufactured in INO containing media using a GMP adaptable process to reach clinically relevant doses that meet standard release criteria and that such cells demonstrate enhanced functionality compared to those manufactured using standard glucose-based media.

## DISCUSSION

Ado is a potent immunosuppressant in the TME and substantial efforts are underway to inhibit Ado to enhance antitumor immune responses[41]. CD39 mediates an early step in Ado generation and is expressed by regulatory T cells[2, 24] and exhausted tumor-reactive T cells in the TME[3–5]. Here, we interrogated the biology of CD39 and Ado in the context of T cell exhaustion and observed that CD39 marks a highly dysfunctional subset of exhausted CD8^+^ T cells that commonly express CD73, manifest phenotypic characteristics of suppressive T cells, and mediate immune suppression via Ado production and signaling via A2aR (**Figure 1A-J, Figure S1A-B**). Knockout of CD39, CD73 or A2aR resulted in modest changes in the phenotype of exhausted CAR T cells.

As an alternative approach, we overexpressed Ada deaminase (ADA), the enzyme that catalyzes Ado into INO, and observed significantly increased frequency of stem cell memory subsets and diminished progenitor exhausted and exhausted subsets, similar to findings reported with overexpression of collagen binding ADA[41, 42]. ADA-OE, but not CD39-, CD73- or A2aR-KO, also induced transcriptional reprogramming that augmented expression of stemness associated genes (**Figure 2B-E, Figure S2D-F**). These results led us to hypothesize that INO, the metabolically active product of ADA catabolism of Ado could reprogram chronically activated T cells toward stemness. INO is a multifunctional intra- and extracellular purine that serves as a carbon source for bioenergetics in human T cells[30], is integrated into tRNA where it plays an important role in gene translation[43] and mediates signals via A2aR and potentially other adenosine receptors. *Bifidobacterium pseudolongum* derived INO augments the effectiveness of anti-CTLA4 therapy in humans and mice via A2aR and IFNγ-dependent induction of Th1 differentiation[44]. In murine models where tumors are unable to utilize INO for bioenergetics, systemic INO augmented the efficacy of anti-CTLA4 and GD2-CAR therapy[30]. In addition to these reported immune activating effects, dendritic cell mediated generation and release of INO via a FAMIN dependent pathway prevented autoimmunity and autoinflammation in murine models[45]. Thus, there is accumulating evidence for immune activating and immunomodulatory effects of INO but mechanistic understanding of its effects in T cells has remained difficult to pinpoint given its pleiotropy.

Our data demonstrates that INO dramatically alters the transcriptome of human T cells to express gene programs associated with stemness (**Figure 3B-C**), decreases expression of FOXP3 and genes involved in purinergic signaling (CD39, CD72 and A2aR), increases protein expression of CD62L, CCR7, CD127 and decreases CD45RA expression, and significantly enhances the function of CAR T cells (**Figure 3J-L**). Metabolic pathways can play a central role in programming T cell differentiation and function[46, 47] and can be regulated by nutrient availability[48, 49]. Consistent with this model, INO exposure diminished T cell proliferation, and reprogrammed the metabolome toward diminished glycolysis, increased glutaminolysis, polyamine synthesis, TCA and oxidative phosphorylation (**Figure 4A-D**). While some effects of INO, such as reduced glycolysis and diminished proliferation, overlap with effects seen with transient glucose restriction[31], the stemness programming effects of INO are distinct and INO induced stemness and metabolic programming was observed regardless of whether glucose was absent or present at a range of physiologic levels (5-18mM). The most notable effects of INO in this system was augmentation of polyamine synthesis, with very significant induction of ornithine decarboxylase, the rate limiting enzyme in generating spermatine, SRM, and EIF5a. Glutaminolysis was also increased by INO and can also drive induction of polyamines synthesis[50]. Increased polyamine levels have been implicated in regulating the TCA cycle via EIF5A hypusination[51] and in inducing chromatin remodeling during Th1 differentiation in mice[34]. INO induced metabolic reprogramming drove major changes in the chromatin accessibility, increasing availability of TF motifs associated with T cell memory and diminishing accessibility of motifs associated with T cell activation (**Figure 4E-H**) Together, the data support a model wherein INO induced metabolic programming drives polyamine synthesis, which in turn augments oxidative phosphorylation and leads to epigenetic programming of T cells toward stemness (**Figure 5G**).

These fundamental observations were readily translated into a clinical scale platform for CAR T cell manufacturing. Using GD2.BBz-CAR T cells, which have shown activity in patients with diffuse midline glioma[11] but whose function has been limited by tonic signaling and early exhaustion[12, 14], we show feasibility of manufacturing CAR T cells in INO containing media and demonstrate enhanced potency of INO manufactured T cells (**Figure 5, S5**). Although the magnitude of T cell expansion was diminished compared to control cultures, sufficient cell numbers were manufactured in 7 days to meet clinical dosing guidelines. GD2-CAR T cells manufactured in INO manifested a more memory-like phenotype, with higher frequencies of naïve and memory populations compared to control, and increased effector function *in vitro* and *in vivo*. In summary, these data demonstrate that INO is a potent inducer of T cell stemness that dramatically modulates T cell metabolism, augments function and induces epigenetic programming. INO can be used to manufacture cell products with enhanced potency compared to standard glucose-based media. Finally, while many approaches are under development to inhibit Ado generation or signaling, our data suggest that approaches designed to metabolize Ado in the TME and generate INO may prove to be more effective in augmenting antitumor immunity.

## MATERIALS AND METHODS

### Viral vector construction

MSGV retroviral vectors encoding the following CARs were previously described: CD19-28z, CD19-BBz, GD2-BBz and Her2-BBz. HA-28z CAR was created as previously described in [14] by introduction of a point mutation into the 14G2a scFv of the GD2-28z CAR plasmid to create the E101K mutation.

### T cell isolation

Healthy donor buffy coats were purchased from the Stanford Blood Center under an IRB-exempt protocol. Primary human T cells were isolated using the RosetteSep Human T cell Enrichment kit (Stem Cell Technologies) according to the manufacturer’s protocol. Isolated T cells were cryopreserved in CryoStor CS10 cryopreservation medium (Stem Cell Technologies).

CD39- T cells were purified using anti-PE MicroBeads (Miltenyi Biotec) and LD autoMACS (Miltenyi Biotec) columns according to the manufacturer’s protocol. Depletion efficiency was assessed by flow cytometry.

### Human CAR T cell production

Non-tissue culture treated 12-well plates were coated overnight at 4°C with 1 ml Retronectin (Takara) at 25 μg/ml in PBS. Plates were washed with PBS and blocked with 2% BSA for 15 min. Thawed retroviral supernatant was added at approximately 1 ml per well and centrifuged for 2h at 32 °C at 3,200 rpm before the addition of cells. Primary human T cells were thawed and activated with Human T-Expander CD3/CD28 Dynabeads (Gibco) at a 3:1 bead:cell ratio in complete medium (RPMI 1640 supplemented with 10% fetal bovine serum, 10 mM N-2- hydroxyethylpiperazine-N9-2-ethanesulfonic acid, 2 mM GlutaMAX, 100 U/mL penicillin (Gibco), and 100 U/mL of IL-2 (Peprotech). T cells were transduced with retroviral vector on day 2 post-activation. Beads were taken off at day 4 post-activation. In INO condition, after removing activation beads at day 4, CAR T cells were centrifugated and resuspended in complete RPMI containing 11mM inosine and no glucose. ADA overexpressing CAR T cells were produced by co-transduction with CAR vector at day 2 post-activation. As surrogate for detection of overexpressed ADA protein, we used anti-HA.11 Epitope Tag antibody (BioLegend).

Clinical-grade GD2.bbz-CAR T cell were manufactured using G-Rex platform (Wilson-Wolf). Apheresis of healthy donors was freshly collected and CD4^+^ and CD8^+^ T cells were enriched using anti-CD4 and anti-CD8 beads (Miltenyi) and EasySep250 (StemCell). T cell activation was performed using Human T-Expander CD3/CD28 Dynabeads (Gibco) at a 1.5:1 bead:cell ratio in complete medium for 3 days. T cells were transduced with GD2.bbz retrovector (MOI=10) on day 2 by spinning at 400g 32°C for 2hrs in the presence of 10ug/ml of protamine sulfate (Frezenius- Cabi). Day 3 post-activation, the beads were taken off and cells were further cultured till day 7 in glucose- or inosine-containing complete RPMI. For small scale, T cells were activated using TransAct (Miltenyi). Day 3 post-activation, transact was washed off and cells were further cultured TexMACS (Miltenyi) containing 3% human serum (ACCESS CELL CULTURE LLC) and 12.5 ng/mL of IL-7 (Miltenyi) and IL-15 (Miltenyi).

### Cell lines

The CD19^+^ Nalm6-GL B-ALL cell line was provided by D. Barrett (Barrett 2011). Nalm6-GD2 was created by co-transducing Nalm6-GL with cDNAs for GD2 synthase and GD3 synthase.

All cell lines were cultured in complete media (CM) (RPMI supplemented with 10% FBS, 10 mM HEPES, 2 mM GlutaMAX, 100 U ml−1 penicillin, and 100 μg ml−1 streptomycin (Gibco)). STR DNA profiling of all cell lines is conducted by Genetica Cell Line testing once per year. None of the cell lines used in this study is included in the commonly misidentified cell lines registry.

### Flow cytometry

All flow analysis was performed at day 14/15 post-T cell activation, unless differently indicated in the text. The anti-CD19 CAR idiotype antibody was provided by B. Jena and L.Cooper. The 1A7 anti-14G2a idiotype antibody was obtained from NCI Frederick and University of Texas M.D. Anderson Cancer Center. Her2.bbz CAR was detected using BFP florescent protein integrated into the plasmid or human Her2-Fc recombinant proteins (R&D). The idiotype antibodies and Fc-fusion protein were conjugated in house with Dylight650 antibody labelling kits (Thermo Fisher). T cell surface phenotype was assessed using the following antibodies:

### From BioLegend

CD4-APC-Cy7 (clone OKT4), CD8-PerCp-Cy5.5 (clone SK1), TIM-3-BV510 (clone F38-2E2), CD39-FITC, PE or APC-Cy7 (clone A1), CD3-PacBlue (clone HIT3a), HA.11 Epitope Tag- PE (16B12)

### From eBioscience

PD-1-PE-Cy7 (clone eBio J105), LAG-3-PE (clone 3DS223H), CD45RO-PE- Cy7 (clone UCHL1), CD45-PerCp-Cy5.5 (clone HI30), CCR7-PE (clone 3D12)

### From BD

LAG-3-BV421 (clone T47-530), CD45RA-FITC or BV711 (clone HI100), CD62L- BV605 (clone DREG-56), CD73-PE-Cy7 or BV510 (clone AD2), CD4-BUV395 (clone SK3), CD8-BUV805 (clone SK1).

### Suppression assay

For assessing IL-2 secretion inhibition 5x10^4^ CD19-BBz-CAR T cells in the presence or absence of 5x10^4^ of control, genetically modified bulk HA-28z-CAR (HA-CAR) T cells were cultured with 5x10^4^ Nalm6 tumor cells in 300uL CM in 96-well flat bottom plates for 24 h. CD8^+^ HA-CAR T cells were selected using anti-PE MicroBeads (Miltenyi) according to the manufacturer’s protocol (average CD8^+^ selection 90%). Triplicate wells were plated for each condition. Culture supernatants were collected and analyzed for IL-2 by ELISA (BioLegend).

### Co-culture assays

HA- or CD19-BBz-CAR T cells between day 14-16 post-activation were cultured with 10 or 40uM of CPI444 (CORVUS BIOPHARMA) for 2 to 24 hours before the duration of coculture (unless stated otherwise) with tumor cells or plate-bound idiotype at the concentration 5ug/ml. To stimulate A2a receptor, cells were treated with 0.01-0.1mM of NECA (Torcis). For cytotoxicity assays, approximately 5x10^4^ of tumor cells were co-cultured with CAR T cells at indicated ratio in 200uL CM in 96-well flat bottom plates. Four images per well at 10x zoom were collected at each time point. Tumor cell growth was quantified by measuring total integrated GFP intensity per well using an IncuCyte ZOOM Live-Cell analysis system (Essen Bioscience) every 2-3hr. GFP signal was normalized to the time 0 signal. Cell culture supernatants were collected at 24 hours, and interleukin-2 (IL-2) and interferon-γ concentrations were determined by enzyme-linked immunosorbent assay (BioLegend). Triplicate wells were plated for each condition.

### CRISPR knockout

CRISPR–Cas9 gene knockout was performed by transient Cas9/gRNA (RNP) complex electroporation using the P3 Primary Cell 4D-Nucleofector X Kit S (Lonza). On day 4 of culture, HA-28z CAR T cells were counted, pelleted and resuspended in P3 buffer at 1.5x10^6^–2x10^6^ cells per 18 μl reaction. 3.3ug Alt-R.Sp. Cas9 protein (IDT) and 120pmol chemically modified synthetic sgRNA (Synthego) (6:1 molar ratio gRNA:Cas9) per reaction was pre-complexed for 10 min at room temperature to create ribonucleoprotein complexes (RNP). A 18-μl cell suspension was mixed with RNP and electroporated using the EO-115 protocol in 16-well cuvette strips. Cells were recovered at 37oC for 30 min in 200 μl T cell medium then expanded as described above. Knockdown efficiency was determined using TIDE or flow cytometry if antibody is available. In the all experiments involving CRISPR-Car9 gene knockout, control CAR T cells and ADA-OE cells were electroporated with a gRNA targeting the safe harbor locus AAVS1[52]. The following gRNA target sequences were used:

AAVS1-GGGGCCACTAGGGACAGGAT, ADORA2a-guide1-GUCUGUGGCCAUGCCCAUCA,

ADORA2a-guide2-UACACCGAGGAGCCCAUGAU (average KO efficiency 86.33%± 4.69 s.e.m.)

CD73-guide1-GCGGGCGCCCGCGCGGCUCG,

CD73-guide2- CUAUGUGUCCCCGAGCCGCG (average KO efficiency 47.75%± 10.04 s.e.m.) CD39- UGGCACCCUGGAAGUCAAAG (average KO efficiency 98%± 0.4472 s.e.m.).

To increase purity of the knock out, CD73^+^ cells were depleted using anti-PE MicroBeads (Miltenyi) according to the manufacturer’s protocol.

### Bulk RNA-seq

For bulk RNA isolation, healthy donor T cells were prepared as described above. On day 14 CD39+ and CD39- CD4+ or CD8+ subsets were isolated using a BD FACSAria cell sorter (Stem Cell FACS Core, Stanford University School of Medicine) and total mRNA was isolated using Qiagen RNEasy Plus mini isolation kit. Bulk RNA-seq was performed by BGI America (Cambridge, MA) using the BGISEQ-500 platform, single-end 50-bp read length, at 30x106 reads per sample.

Reads mapping and counting were performed using STAR (version 2.7.6a) with GRCh38 as reference genome, HTSeq-count tools respectively. Genes having less than 10 counts in all samples in total were filtered out. Variance stabilization was applied to raw counts using the rlog() function of DESeq2 (version 1.30.0) prior to unsupervised analysis. For clustering analysis, most variable genes between HA and ADA-OE HA were selected using 2.7 as log2 fold-change threshold, and 1e-15 as adjusted p-value threshold. Euclidean correlation and Ward’s method were used respectively as distance metric and agglomeration criterion. Principal Component Analysis was performed using pcaExplorer tool (version 2.16.0) with default parameters.

DAVID gene annotation enrichment analysis was performed using KEGG pathways and GO terms (biological process, cellular component, and molecular function). Functional annotation clustering was performed and terms with p < 0.05 (Benjamini corrected) are shown. Redundant terms were manually removed for visualization.

### ATP measurements

CAR T cells were washed with RPMI without phenol red (Agillent). Next, 5x10^4^ of cells were resuspended in 150ul of phenol free RPMI media and spiked with 20uM of ATP (PerkinElmer). After 10min of incubation at 37°C supernatants were collected and concentration of ATP/sample was measured using ATPlite Luminescence Assay System (PerkinElmer) according to the manufacturer’s protocol.

### ADO measurements

To measure the ability of ectoenzymes at the surface CAR T cells were washed with RPMI without phenol-free (Agillent). Next, 5x10^4^ of cells were resuspended in 150ul of phenol-free RPMI media and spiked with 20uM of ATP (PerkinElmer). After 30min of incubation at 37°C supernatants were collected. Ado concentration was assessed using Ado Assay Kit (abcam) according to the manufacturer’s protocol.

### Luminex

At day 14 post-activation sorted CD39+ and CD39- CD4+ or CD8+ CAR T cells were co cultured with Nalm6-GD2 cells at 1:1 ratio. Duplicate wells were plated for each condition. After 24hrs supernatants were collected and analyzed using Luminex assay performed at the Human Immune Monitoring Center of Stanford University. Human 62-plex kits were purchased from eBioscience/Affymetrix and used according to the manufacturer’s recommendations with modifications as described. Briefly, beads were added to a 96-well plate and washed in a BioTek ELx405 Select Deep Well Washer. Samples were added to the plate containing the mixed antibody-linked beads and incubated at room temperature for 1 h followed by overnight incubation at 4 °C with shaking. The cold and room temperature incubation steps were performed on an orbital shaker at 500–600 r.p.m. After overnight incubation, plates were washed in a BioTek ELx405 Select Deep Well Washer; then, biotinylated detection antibody was added for 75 min at room temperature with shaking. The plate was washed as described earlier and streptavidin-PE was added. After incubation for 30 min at room temperature, washing was performed as described earlier and reading buffer was added to the wells. Plates were read using a Luminex FLEXMAP 3D instrument with a lower bound of 50 beads per sample per cytokine. Custom assay control beads by Radix Biosolutions were added to all wells. The dilution factor was accounted for.

For each cytokine, concentration [pg/ml] was calculated.

### QuantiBRITE antigen density quantification assay

The cell surface quantification of GD2 on tumor lines was enumerated by flow cytometry using BD QuantiBRITE beads and Custom Quantitation Beads (BD Biosciences, San Jose, CA, USA) according to the manufacturer’s protocol.

### Mass cytometry

1×10^6^ cells were washed with two times with PBS (Rockland) and then resuspended in 1ml of 250nM cisplatin and PBS (Fluidigm) for assessing cell viability. Cells were incubated for 3min at RT and washed with cell staining medium (CSM, 1X PBS with 0.05% BSA, 0.02% sodium azide). Then cells were fixed with 1.6% paraformaldehyde diluted in PBS for 10 min at room temperature, followed by a wash with 1X PBS. Samples were subsequently frozen using CryoStore (Thermo Fisher Scientific) for further use. Upon thawing and washing in CSM, barcoding was performed using a palladium-based approach using the Cell-ID 20-Plex Pd Barcoding Kit manufacturer’s protocol (Fluidigm Corporation). Cell samples were mixed and treated as one sample for all subsequent steps. Titrated amounts of each cell surface antibody were mixed, filtered through a 0.1 mm spin filter (Millipore), and added to the merged barcoded sample for 30 min at room temperature. Cells were then washed two times with CSM and permeabilized by adding ice cold methanol for 10 min at 4°C, followed by two washes with CSM. Finally, samples were resuspended in IR- intercalator (0.5 mM iridium intercalator and 1% PFA in 1X PBS), washed one time with CSM and two times with ddH2O, and filtered through a 50 mm cell strainer (Thermo Fisher). Cells were then resuspended at 1 × 10^6^ cells/ml in ddH2O with 1x EQ four-element beads (Fluidigm Corporation, no. 201078). Cells were acquired on a Fluidigm Helios mass cytometer. Data was analyzed using OMIQ software.

FlowSOM analysis was performed using: TCF7, CD27, Cd278, CD69, CD62L, CD28, CD45RO, CD244, CD38, TIM3, TIGIT, CD137, GLUT1, T-BET, CD25, CD272, CD223, cJUN, CCR7, CD127, CD45RA, Ki67, CD122, PD1, TOX and LEF1 markers.

### ATAC-seq

Approximately 100k CAR-T cells were washed in ice-cold PBS and subjected for nuclei isolation using the following lysis buffer: 10mM Tris-HCl pH 7.5, 10mM NaCl, 3mM MgCl2, 0.1% Tween- 20, 0.1% NP40, 0.01% Digitonin and 1% BSA. After washing the cells, 50ul lysis buffer was added to each sample and cells were resuspended by pipetting. Nuclear pellets were centrifuged and resuspended in the transposase reaction containing 10.5ul H2O, 12.5ul 2xTD buffer and 2ul Tn5 transposase in total of 25ul. The reaction was incubated for 30 minutes at 37C. Reaction was stopped by the addition of 75ul TE buffer and 500ul PB buffer (Qiagen), followed by column purification per manufacturer’s recommendation (Qiagen, Minelute Kit). DNA was eluted from the columns in 22ul H2O. PCR reactions were set up as follows: 21ul DNA, 25ul Phusion master mix (NEB) and 2ul of each barcoded PCR primer (ApexBio, K1058). 14 PCR cycles were run for each sample. Reactions were clean up with AMPure XP beads according to the recommendations of the manufacturer. Libraries were quantified with Qubit fluorometer and fragment analysis was performed with Bioanalyzer. Libraries were sequenced on a NovaSeq 6000 sequencer.

### ATAC-Seq data preprocessing and analysis

The ENCODE ATAC-seq pipeline v2.0.3 was used for quality control and processing of ATAC-seq using default parameters. Briefly, raw ATAC-seq reads were trimmed with cutadapt to remove Illumina Nextera adapter sequences and aligned to hg19 using Bowtie2. Uniquely aligned reads were identified using picard MarkDuplicates with 73–85% non-duplicates mapped reads among all the samples. Peak from individual samples was called on Tn5-corrected insertion sites using MACS2. Reads from mitochondria and reads of low mapping quality (MAPQ < 30) were excluded. A union peak set was compiled by extending peak summits to 500 bp, merging all summits, running bedtools cluster, selecting summits with the highest MACS2 score. Peaks intersecting with ENCODE hg19 blacklisted regions or regions on chrY and chrX were removed. For visualization, consensus peaks from biological replicates were merged and genome coverage plot was generated by IGV v1.9.0[53]. Only peaks with at least 5 counts-per-million in at least one of the samples were included in the downstream analyses. PCA was performed on the 5000 most variable peaks using the regularized log transform values from DESeq2 v1.30.1[54]. For differential peak accessibility analysis and transcription factor motif analysis batch effects by donor were corrected with the function ComBat_seq from the sva package v3.38.0[55]. Transcription factor motif deviation analysis was carried out using chromVAR v1.12.0 as previously described[56] with human motifs from the JASPAR database. Motifs with significant differences (adjusted p-value <0.05) in deviations between in HA-INO and HA were ranked based on the size of the deviation difference. DESeq2 was used to identify peaks with significant (adjusted p-value < 0.001) differential accessibility, of which two peak sets were established: one containing the 500 peaks with the strongest decrease and one with the 500 peaks with the strongest increase in accessibility in HA-INO versus HA. For these two peak sets relative motif enrichment was computed with HOMER’s findMotifsGenome.pl using default parameters. The transcription factor motifs were ranked based on the p-values of the enrichment level.

### Mice

Immunocompromised NOD-SCID-Il2rg−/− (NSG) mice were purchased from JAX and bred in-house. All mice were bred, housed, and treated in ethical compliance with Stanford University IACUC (APLAC) approved protocols. Six-to-eight-week-old male or female mice were inoculated with either 1 × 10^6^ Nalm6-GL leukemia via intravenous or 1 × 10^6^ 143B osteosarcoma via intramuscular injections. All CAR T cells were injected intravenously and media swap to inosine-supplemented media was performed at day 6 post activation. Time and treatment dose are indicated in the figure legends. Bioluminescence imaging was performed using a Spectrum IVIS instrument. Values were analyzed using Living Image software. Solid tumor progression was followed using caliper measurements of the injected leg area. Mice were humanely euthanized when an IACUC-approved end-point measurement reached 1.75 cm in either direction (for solid tumor) or when mice demonstrated signs of morbidity and/or hind-limb paralysis (leukemia). Mice were randomized to ensure equal pre-treatment tumor burden before CAR T cell treatment.

### Blood analysis

Blood was collected from the retro-orbitalsinus into Microvette blood collection tubeswith EDTA (Fisher Scientific). Whole blood was labeled with anti-CD45 (HI30, eBioscience) or anti-CD3 (HIT3a, BioLegend). Next, red blood cells were lysed with FACS Lysing Solution (BD) according to manufacturer’s instructions. Samples were mixed with CountBright Absolute Counting beads (ThermoFisher) before flow cytometry analysis.

### Statistical analysis and graphical design

Unless otherwise noted, statistical analyses for significant differences between groups were conducted using unpaired two-tailed t-tests without correction for multiple comparisons and without assuming consistent s.d. using GraphPad Prism 8.4. Graphical abstract and schematics were design in biorender.com.

### Data availability

The main data supporting the findings of this study are available within the article and its Supplementary information. All raw data generated during the study are available from the corresponding authors upon reasonable request

## Author contributions

C.L.M. supervised the study and acquired funding. D.D.K., E.S. and C.L.M. designed and conceptualized the study. D.D.K cloned the construct, designed and performed experiments, and analyzed data. M.M, L.S., C.F., N.G., J.M-A., M.B., B.D., K.S. performed experiments and collected the data. E.S, K.M, S.M., A.L. performed biostatistical analysis. V.K., J.A-G., C.F., P.X., J.H. performed animal experiments and collected the data. A.T.S., S.P., S.F. advised on the experimental design. D.K., E.S., and C.L.M. wrote the manuscript. All other authors commented on and revised the manuscript.

Supplementary 1. CD39 expression depends on tonic signaling and is crucial for generation of immunosuppressive adenosine.

A. (Top) Volcano plots of RNA expression levels in CD39^−^ vs. CD39^+^ HA-CAR T cells at day 14 post-activation. Significantly different genes identified by DESeq2 (Wald test) are shown in red

(padj < 0.1). (Bottom) Venn diagram depicting overlap between genes significantly upregulated in CD39^+^ vs. CD39^−^ CD4^+^ and CD8^+^ T cells.

B. Gene Set Enrichment Analysis (GSEA) of bulk RNA-seq collected 14 days post-activation of CD39^+^compared to CD39^−^HA-CAR T cells using dataset from the indicated publications.

C. Day 16 post-activation unselected or sorted CD39^−^HA-CAR T cells were cultured with dasatinib (1uM) or neutralizing anti-TGFβ (0.001mg/ml). CD39 expression was measured 7 days later by flow cytometry. (Left) Contour plots from a representative donor and (right) mean fold change ± s.e.m. of CD39 expression from n=4 donors shown. *, *P* < 0.05; **, *P* < 0.01; ***, *P* < 0.001; ****, *P* < 0.0001

D. Percent of ATP hydrolyzed by mock or CAR-expressing T cells with or without CD39 or CD73 after spiking with 20uM of ATP at day 15 post-activation. Representative data from n=4 donors. Average purity of the knock-out cells was > 90%. P values determined by unpaired two-tailed t- tests. *, *P* < 0.05; **, *P* < 0.01; ***, *P* < 0.001; ****, *P* < 0.0001

E. IL-2 secreted by HA- or CD19.bbz-CAR T cells stimulated with 1A7 anti-idiotype mAb (5 ug/ml)

± NECA (0.01mM) and A2aR inhibitor (CPI444, 40 uM) at day 16 post-activaiton. Data are mean ± s.e.m. from triplicate wells. Representative data of n= 2-3 donors shown. P values determined by unpaired two-tailed t-tests. *, *P* < 0.05; **, *P* < 0.01; ***, *P* < 0.001; ****, *P* < 0.0001

F. At day 16 post-activation CD19.bbz-CAR T cells were activated with Nalm6 alone ± HA-CAR or enriched CD8^+^ HA-CAR T cells at 1:1 ratio. IL-2 secretion was assessed 24hrs post-stimulation. Data are mean ± s.e.m. from triplicate wells. Representative experiment of total n=2 is shown.

*, *P* < 0.05; **, *P* < 0.01; ***, *P* < 0.001; ****, *P* < 0.0001

Supplementary 2. ADA-OE decreases CAR T cell exhaustion.

A. (Left) Design of the ADA vector and (right) cell surface expression of ADA detected by staining for ha-tag at D10 and measured by flow cytometry. SS- leader/signaling domain; ha-tag- hemagglutinin tag; ADA- adenosine deaminase; CD8-TM- CD8 transmembrane sequence.

B. Representative surface expression of the CAR receptor by control HA-CAR vs. ADA-OE HA-CAR T cells at day 10 post-activation.

C. GSEA of day 15 ADA-OE vs. HA-CAR T cells using dataset from the indicated publications.

D. CyTOF analysis of HA-CAR T cells at day 15 post-activation. Graphs represent median expression of indicated markers by CD8^+^ CAR T cells. Total of n=4 donors shown. P values determined by paired two-tailed t-tests. *, *P* < 0.05; **, *P* < 0.01; ***, *P* < 0.001; ****, *P* < 0.0001

E-F. UMAP analysis of day 15 CD4^+^ HA-CAR cells engineered as in (Figure 2A). Expression of 26 markers was analyzed by CyTOF. 5,000 or maximum of CD4^+^ CAR T cells from each donor (n=4) were combined and colored by (E) genotype or (F) subpopulation defined by FlowSOM algorithm. Pie charts show population frequencies defined using FlowSOM for each condition. Table shows the average frequencies from n=4 donors and corresponding p values determined by paired two-tailed t-tests.

Supplementary 3. HA-INO-CAR T cells exhibit decreased CD73 expression and increased effector function in vitro

A. (Left) Representative contour plots of CD39 and CD73 expression by HA-CAR T cells measured by flow cytometry at day 14/15 post-activation. (Right) Percentage of CD39^+^, CD73^+^ or double positive CD39^+^CD73^+^ T cells in CD8^+^ HA-CAR T cells in 7 donors from independent experiments. P values determined by paired two-tailed t-tests. *, *P* < 0.05; **, *P* < 0.01; ***, *P* < 0.001; ****, *P* < 0.0001

B. Flow cytometry analysis of CD62L expression by CD4^+^ HA-CAR T cells at day 21 post-activation. Histograms of representative donor (Left) and pooled data from n=8 donors shown (Right). P values determined by paired two-tailed t-tests. *, *P* < 0.05; **, *P* < 0.01; ***, *P* < 0.001; ****, *P* < 0.0001

C. UMAP plots of day 14 CD8^+^ (Top) and CD4^+^ (Bottom) HA- or HA-INO-CAR T cells analyzed using CyTOF. 9,000 or maximum of CD4^+^ CAR T cells from each condition from n=4 donors were organized by their combined expression of 5 exhaustion markers. Graphs represent median expression of indicated markers. P values determined by paired two-tailed t-tests. *, *P* < 0.05; **, *P* < 0.01; ***, *P* < 0.001; ****, *P* < 0.0001

D. Histograms showing GD2 antigen expressed by Nalm6-GD2, mg63.3 and 143b tumor lines with corresponding number of molecules per cell.

E. IFNγ secretion by day 14 HA-CAR T cells and HA-INO-CAR T cells, 24hrs post-stimulation with Nalm6-GD2 (left) or 143b (right) tumor lines. Error bars represent mean ± SD of triplicate wells from one representative donor of n=1-2 donors. P values determined by unpaired two-tailed t-tests.

*, *P* < 0.05; **, *P* < 0.01; ***, *P* < 0.001; ****, *P* < 0.0001

F. Co-cultures of HA, HA-ADA-O/E- or HA grown in INO stimulated with Nalm6-GD2 (1:8 E:T) at day 14 post-activation. Tumor GFP fluorescence intensity was normalized to the first time point (duplicate or triplicate wells). Representative donor of n=2 donor shown. P values determined by two-way ANOVA with Dunnett’s multiple comparisons test. *, *P* < 0.05; **, *P* < 0.01; ***, *P* < 0.001; ****, *P* < 0.0001

G. Schematic of the experimental design.

H. Relative frequency of terminally differentiated effector (TEMRA; CD45RA^+^CD62L^−^), stem cell memory (CD45RA^+^CD62L^+^), central memory (CM; CD45RA^−^CD62L^+^), and effector memory (EM; CD45RA^−^CD62L^−^) in CD8^+^ HA-CAR T cells cultured in different media conditions at day 21 post-activation measured by flow cytometry. (Top) Dot plots of representative donor shown. (Bottom) Frequencies of different populations polled from n=5-9 donors from independent experiments. P values determined by paired two-tailed t-tests *, *P* < 0.05; **, *P* < 0.01; ***, *P* < 0.001; ****, *P* < 0.0001

Supplementary 4. INO improves exhausted and non-exhausted CAR T cells function in vivo

A. NSG mice were inoculated with 1 × 10^6^ Nalm6-GD2 leukemia cells. On day 3, 0.5 × 10^6^ or 2 × 10^6^ number of HA-CAR^+^ T cells or mock cells were transferred intravenously. Tumor growth was monitored by bioluminescent imaging. Data are mean ± s.e.m. of n=5 mice per group. P values determined at day 18 by using Mann-Whitney test. *, *P* < 0.05; **, *P* < 0.01; ***, *P* < 0.001; ****, *P* < 0.0001

B. NSG mice were inoculated with 1 × 10^6^ Nalm6-GD2 leukemia cells. On day 3, 0.5 × 10^6^ number of HA-CAR^+^ T cells or 2 × 10^6^ number of mock cells were transferred intravenously. Long term survival of CAR-treated mice. Survival curves were compared using the log-rank Mantel–Cox test.

C. IFNγ secretion by CD19.28z-CAR T cells 24hrs post-stimulation with Nalm6 tumor line expressing endogenous or low CD19 antigen density. Representative donor shown of n=2. Error bars represent mean ± SD of triplicate wells from one representative donor (n=2 donors). P values determined by unpaired two-tailed t-tests. *, *P* < 0.05; **, *P* < 0.01; ***, *P* < 0.001; ****, *P* < 0.0001

D. NSG mice were injected intravenously with 1 × 10^6^ Nalm6 leukemia cells, and then 0.2 × 10^6^ mock, CD19.28z-CAR T cells manufactured in the presence of inosine or in regular media were given intravenously on day 3. Tumor progression was monitored using bioluminescent imaging.

Data are mean ± s.e.m. of n = 5 mice per group from one experiment. P values determined at day 18 by Mann-Whitney test. *, *P* < 0.05; **, *P* < 0.01; ***, *P* < 0.001; ****, *P* < 0.0001

E. NSG mice were inoculated with 1 × 10^6^ 143B osteosarcoma cells via intramuscular injection. At D4 post-engraftment 1 × 10^7^ mock or Her2.bbz-CAR T cells manufactured in glucose- or inosine-containing media were given intravenously. Pooled measurements from two independent experiments shown. *, *P* < 0.05; **, *P* < 0.01; ***, *P* < 0.001; ****, *P* < 0.0001

Supplementary 5. Addition of inosine into GMP-grade TexMACS culture media increases CAR T cells function in vitro.

A. Schematic of the experimental design.

B. Viability of GD2.bbz-CAR T cells cultured for 4 days in TexMACS containing indicated concentration of inosine at day 7 post-activation. N=8 donors from independent experiments. P values determined by paired two-tailed t-tests. *, *P* < 0.05; **, *P* < 0.01; ***, *P* < 0.001; ****, *P* < 0.0001

C. Expansion rate between day 3 and day 7 post-activation of CAR T cells grown in TexMACS containing indicated concentration of inosine. N=8 donors from independent experiments. P values determined by paired two-tailed t-tests. *, *P* < 0.05; **, *P* < 0.01; ***, *P* < 0.001; ****, *P* < 0.0001

D. Percentage of CD62L^+^CD8^+^ GD2.bbz-CAR T cells at day 10 post-activation grown in TexMACS containing indicated concentration of inosine. N=6 donors from independent experiments. P values determined by paired two-tailed t-tests. *, *P* < 0.05; **, *P* < 0.01; ***, *P* < 0.001; ****, *P* < 0.0001

E. GD2.bbz-CAR T cells grown in media containing indicated inosine concentration between day 3 and day 10 post-activation and stimulated with 143b tumor. IL-2 and IFNγ secretion after 24hrs of co-culture was measured. Data are mean ± s.d. of duplicate or triplicate wells. Representative of n=3 donors shown. P values determined by unpaired two-tailed t-tests. *, *P* < 0.05; **, *P* < 0.01; ***, *P* < 0.001; ****, *P* < 0.0001

