## supplementary figures for "Inosine Induces Stemness Features in CAR T cells and Enhances Potency"

Supplementary 1

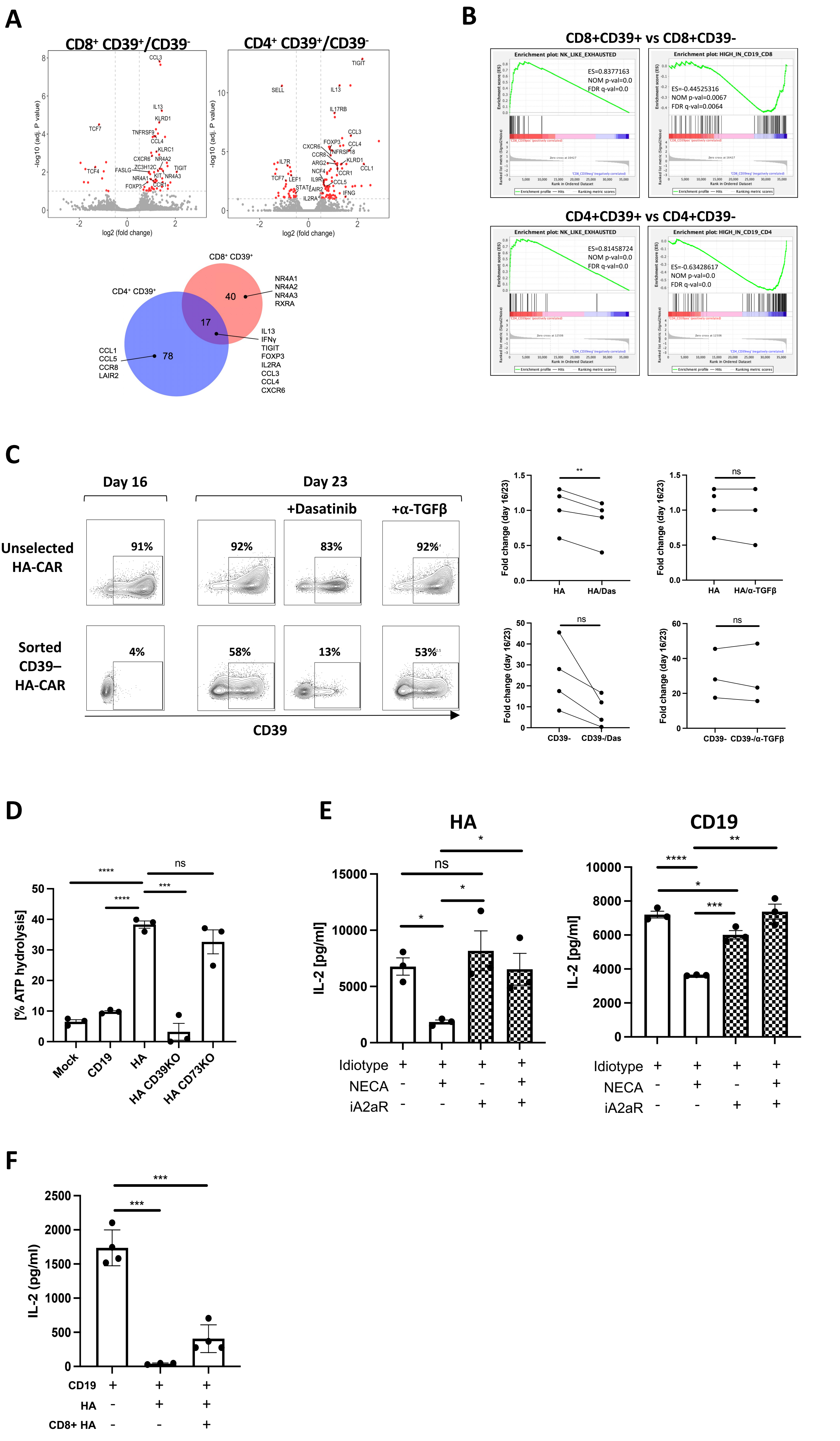

### Supplementary 2

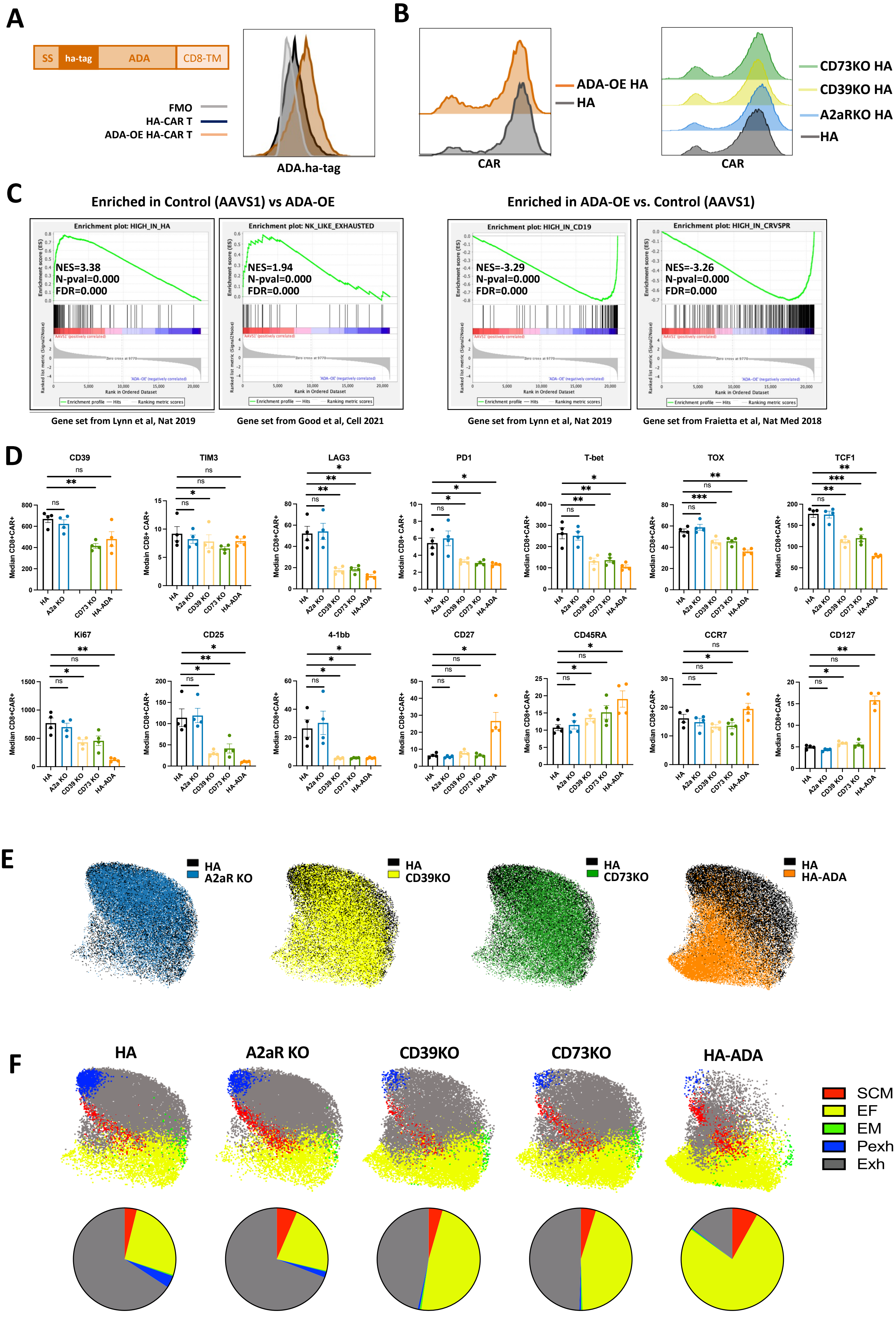


| CD4+ | HA | A2aR KO | | CD39 KO | | CD73 KO | | ADA-OE | |
|---|---|---|---|---|---|---|---|---|---|
| | Frequency (%) ±s.e.m. | Frequency (%) ±s.e.m. | P-value | Frequency (%) ±s.e.m. | P-value | Frequency (%) ±s.e.m. | P-value | Frequency (%) ±s.e.m. | P-value |
| SCM | 3.87±2.574 | 6.47±2.682 | ns | 4.41±1.902 | ns | 4.81±2.441 | ns | 8.11±1.728 | 0.0250 |
| EF | 26.1±6.45 | 22.2±10.392 | ns | 47.88±13.961 | 0.0187 | 44.65±12.961 | ns | 76.78±6.782 | 0.0032 |
| EM | 0.41±0.145 | 0.28±0.118 | ns | 0.55±0.154 | ns | 0.47±0.121 | ns | 0.42±0.084 | ns |
| Pexh | 3.83±2.147 | 1.9±0.73 | ns | 0.59±0.465 | 0.0321 | 0.53±0.144 | ns | 0.27±0.053 | 0.0471 |
| Exh | 65.8±4.323 | 69.18±8.533 | ns | 46.58±13.182 | 0.0321 | 49.53±12.613 | ns | 14.41±5.255 | 0.0013 |

Supplementary 3

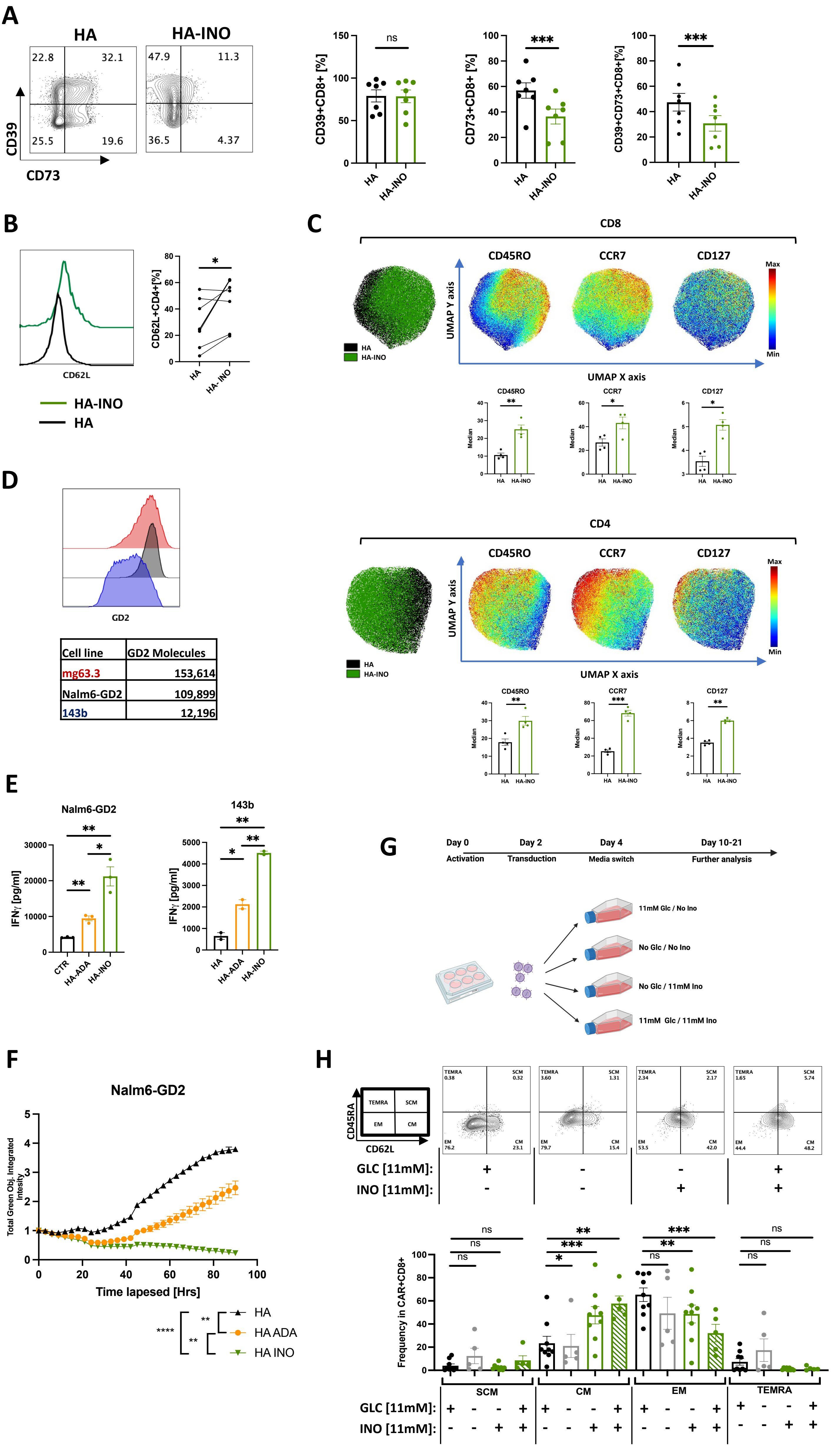


| Cell line | GD2 Molecules |
|---|---|
| mg63.3 | 153,614 |
| Nalm6-GD2 | 109,899 |
| 143b | 12,196 |

#### Supplementary 4

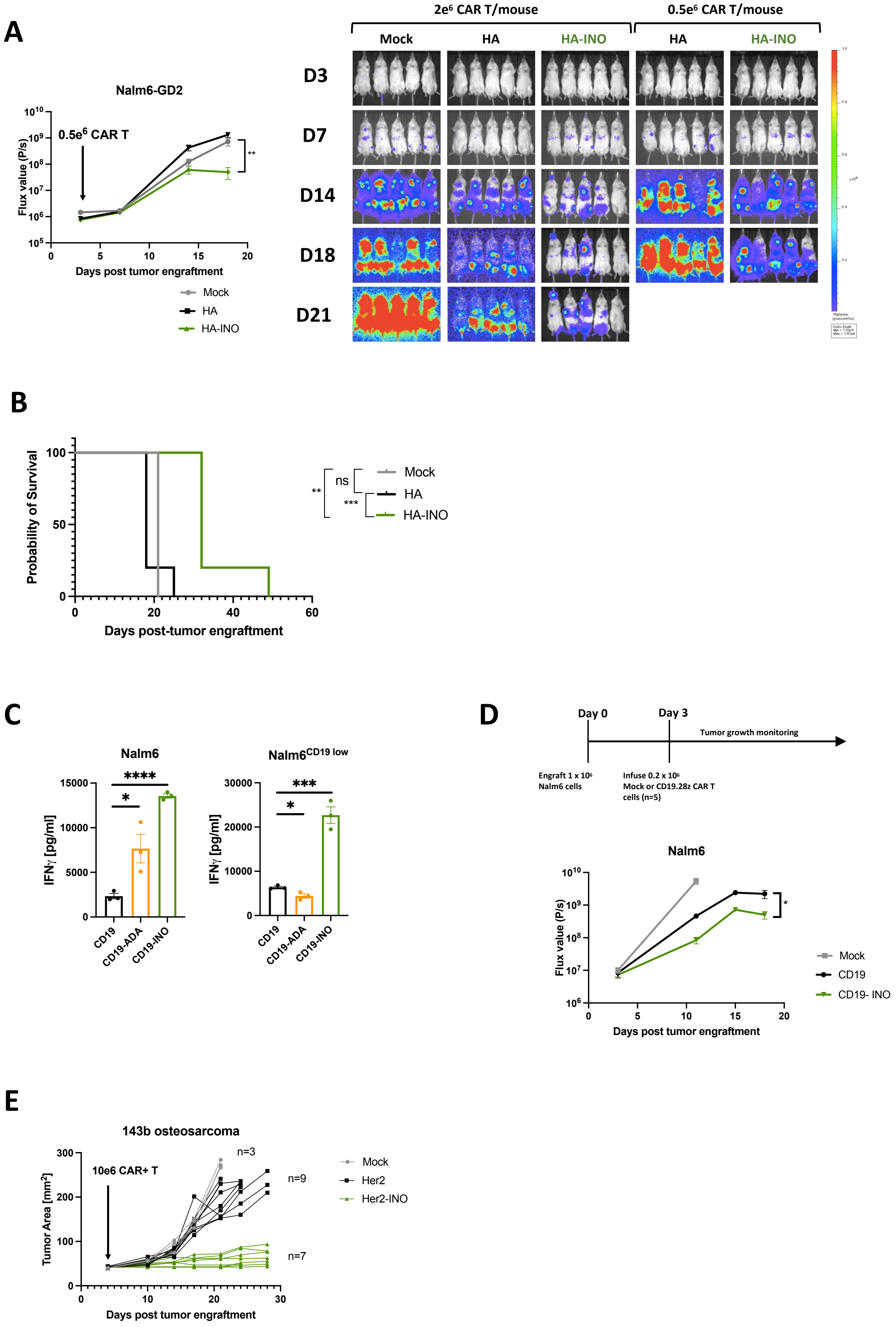

### Supplementary 5

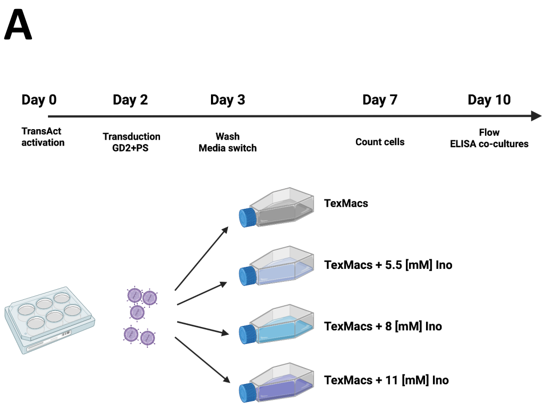


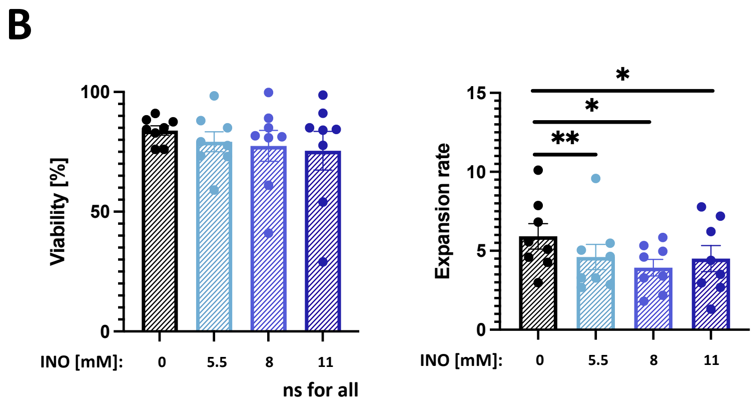


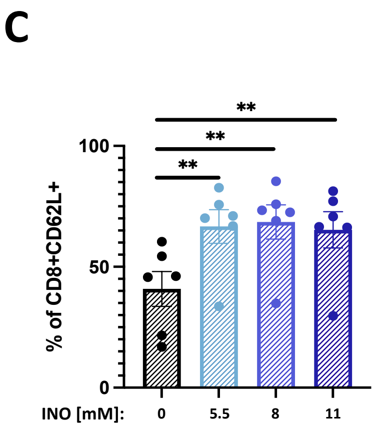


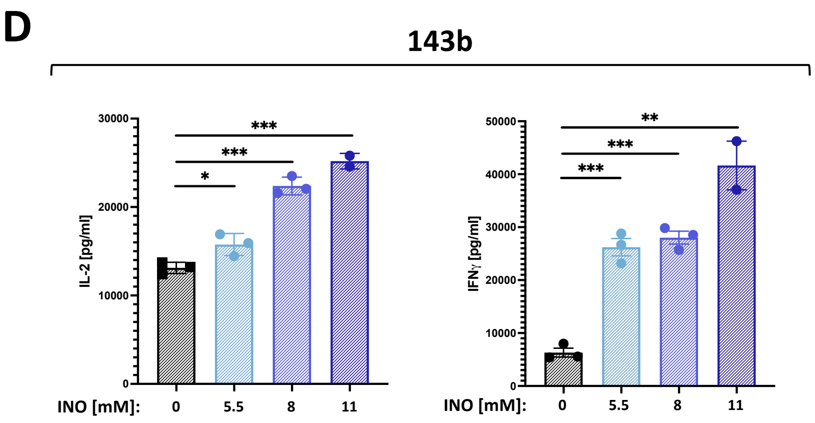
